# Exploration of the main active metabolites from *Tinospora crispa* (L.) Hook. f. & Thomson stem as insulin sensitizer in L6.C11 skeletal muscle cell by integrating *in vitro*, metabolomics, and molecular docking

**DOI:** 10.1101/2023.09.01.555830

**Authors:** Ummu Mastna Zuhri, Nancy Dewi Yuliana, Fadilah Fadilah, Linda Erlina, Erni Hernawati Purwaningsih, Alfi Khatib

## Abstract

**Ethnopharmacological relevance:** *Tinospora crispa* (L.) Hook. f. & Thomson stem (TCS) has long been used as folk medicine for the treatment of diabetes mellitus. Previous study revealed that TCS possesses multi-ingredients and multi-targets characteristic potential as insulin sensitizer activity. However, its mechanisms of action and molecular targets are still obscure.

**Aim of the study:** In the present study, we investigated the effects of TCS against insulin resistance in muscle cells through integrating *in vitro* experiment and identifying its active biomarker using metabolomics and in molecular docking validation.

**Materials and methods:** We used centrifugal partition chromatography (CPC) to isolate 33 fractions from methanolic extract of TCS, and then used UHPLC-Orbitrap-HRMS to identify the detectable metabolites in each fraction. We assessed the insulin sensitization activity of each fraction using enzyme-linked immunosorbent assay (ELISA), and then used confocal immunocytochemistry microscopy to measure the translocation of glucose transporter 4 (GLUT4) to the cell membrane. The identified active metabolites were further simulated for its molecular docking interaction using Autodock Tools.

**Results:** The polar fractions of TCS significantly increased insulin sensitivity, as measured by the inhibition of phosphorylated insulin receptor substrate-1 (pIRS1) at serine-312 residue (ser312) also the increasing number of translocated GLUT4 and glycogen content. We identified 58 metabolites of TCS, including glycosides, flavonoids, alkaloids, coumarins, and nucleotides groups. The metabolomics and molecular docking simulations showed the presence of minor metabolites consisting of tinoscorside D, higenamine, and tinoscorside A as the active compounds.

**Conclusions:** Our findings suggest that TCS is a promising new treatment for insulin resistance and the identification of the active metabolites in TCS could lead to the development of new drugs therapies for diabetes that target these pathways.

**Highlights:**

TCS attenuated insulin resistance in skeletal muscle through lowering IRS ser312 phosphorylation, increasing translocation of GLUT4 and glycogen content.
Tinoscorside D, tinoscorside A, and higenamine were identified as the primary bioactive compounds of TCS.

## 1. Introduction

*Tinospora crispa* (L.) Hook. f. & Thomson is a medicinal plant that belongs to the genus *Tinospora* of the *Menispermaceae* family. It is found in primary rainforests or mixed deciduous forests of Southeast Asia (Indonesia, Thailand, and Malaysia), Africa, and the northeastern region of India (Ahmad et al., 2016). *T. crispa* has a variety of local names that follow the language of the region where it grows, including Brotowali, Antawali, and Andawali (Indonesia); Petawali and Akar Seruntum (Malaysia); Giloya (India); Makabuhay, Panyawan, and Manunggal (Philippines) (Heyne, 1987; Keim & Sujarwo, 2021). The synonym names were diverse, including *Menispermum crispum* L.; *Menispermum rimosum* (Blume) Spreng.; *Menispermum tuberculatum* Lam.; *Menispermum verrucosum* Roxb.; *Tinospora rumphii* Boerl.; *Tinospora tuberculata* (Lam.) Beumée ex K. Heyne; *Tinospora verrucosa* (Roxb.) W.Theob. (Keim & Sujarwo, 2021).

Empirically, it has been commonly used in traditional medicine to treat various health conditions. In Indonesia, since the early 20th century, the decoction of *T. crispa* stems has been empirically reported for the therapy of diabetes mellitus as well as a postpartum remedy and muscle pain treatment (Heyne, 1987; Keim & Sujarwo, 2021). In Thailand, the decoction from the stem of *T. crispa* has been used as an antipyretic, anti-inflammatory, and appetite enhancer (Ahmad et al., 2016). In Malaysia, it is traditionally used for numerous therapeutic purposes such as diabetes, hypertension, appetite stimulation, and protection from mosquito bites (I. Ismail et al., 1931).

Scientific studies have shown that *T. crispa* has a variety of pharmacological activities, including anti-inflammatory, cytotoxic against cancer cell lines, antimalarial, hypoglycaemic, and antifilarial. It also has antioxidant activity, immunomodulatory effect, antinociceptive activity, and hypoglycemic effect (Ahmad et al., 2016; Gao et al., 2016; Hamid et al., 2015; Haque et al., 2020; Klangjareonchai & Roongpisuthipong, 2012; Lam et al., 2012; Lokman et al., 2013; Noor & Ashcroft, 1989; Ruan et al., 2012; Thomas et al., 2016; Zuhri et al., 2022). The phytochemical composition of *T. crispa* includes groups of alkaloids, glycosides, diterpenoids, triterpenoids, lactones, steroids, and flavonoids (Ahmad et al., 2016; Thomas et al., 2016). The diterpenoid glycosides in *T. crispa* include borapetosides A-G, with three main constituents as borapetoside A, B, and C. Among others, borapetoside A has been shown to have the highest activity in lowering plasma glucose levels and increasing plasma insulin levels intraperitoneally (Fukuda et al., 1985, 1986; Lam et al., 2018; Ruan et al., 2013; Yonemitsu et al., 1995). Of the Literature studies indicate that the antidiabetic activity of *T. crispa* is a combination of several mechanisms related to insulin secretion, inhibition of glucose absorption, and insulin signaling in muscle and liver tissues (Ahmad et al., 2016; Thomas et al., 2016; Zuhri et al., 2022).

The commonly used method in natural product-based drug discovery research, known as bioassay-guided isolation, is considered as less accurate in identifying bioactive compound markers from natural sources. This approach is less representative for research on complex natural materials (Yuliana et al., 2013). However, to date, the target for insulin sensitization therapy in skeletal muscle and its corresponding bioactive compounds remain undisclosed. Therefore, there is a need for research on the *T. crispa* using approaches that better represent its activities and mechanisms as a multicomponent system, such as bioinformatics and metabolomics. In this research, we employed this innovative methodology to investigate the pharmacological activity of *T. crispa* stem (TCS) as insulin sensitizer *in vitro* and determine its main active constituents, which were subsequently validated using *in silico* molecular docking study.

## 2. Materials and methods

### 2.1. Reagents and antibodies

TCS was purchased from plantation in Temanggung, Central Java Province, Indonesia. All chemicals were purchased from Sigma-Aldrich (St. Louis, MO, USA) unless stated otherwise. N-hexane, ethyl acetate, methanol, metformin hydrochloride, L6.C11 rat skeletal muscle myoblast cell line (ECACC 92102119, ETACC), fetal bovine serum (FBS, Gibco), Dulbecco’s Modified Eagle Medium (DMEM, ThermoFisher), rat pIRS1 Elisa kit (MBS1605652, MyBioSource), primary antibody GLUT4 conjugated alexa fluor 488 (SC-53566 AF488, Santa Cruz), and rat glycogen ELISA kit (MBS729293, MyBioSource).

### 2.2. Extraction and fractionation of TCS

Dried TCS (2.9 kg) were extracted with 80% MeOH in water (10 L) at room temperature in ultrasonic water bath for 30 minutes. The filtrate than evaporated in a vacuum rotary evaporator and dried by lyophilization under reduced pressure in −103 °C. Crude MeOH dried extract (79.9 g) was obtained and fractionated in CPC using H Arizona solvent system (n hexane-ethyl acetate-methanol-water; 1-3-1-3) in descending mode (Berthod et al., 2005). 33 fractions were collected from triplicate fractionation coded by its eluted order from F1 to F11 and coded with A, B, C after the number for the replication order. Each fraction then evaporated, dried under lyophilization, and was subjected to further analysis.

### 2.3. Cell treatment

L6.C11 skeletal muscle cells were maintained in growth medium, consist of DMEM supplemented with 10% (v/v) fetal bovine serum (FBS), 1% (v/v) 100 units/mL penicillin, 100 mg/mL streptomycin, and 1% (v/v) 250 μg/mL amphotericin B at 37 °C incubator in a humidified 5% CO2 atmosphere. The cells were seeded at 96 well plate (8000 cells/well) for pIRS1, 48 well plate (24000 cells/well) for glycogen measurements while for GLUT4 measurement were seeded at 12 well plate (150000 cells/well) equipped with sterilized round cover slips at the bottom of each plate. After the cells reach 80-90% confluence, the FBS was switched to 2% (v/v) to induce differentiation. It was maintained for 6 days with medium changing every 2 days to fully differentiated myotubes (Tang et al., 2017).

Insulin resistance induction was conducted according to previous experimental method (Huang et al., 2002; Zuhri, 2022). Myotube cells were pretreated with high glucose-high insulin medium (25 mmol/L glucose and 100 nmol/L insulin) for 24 hours followed by deprived of serum and glucose (serum free and 5 mmol/L glucose) for 5 hours before incubation with tested materials. The tested fractions were 400 ppm of 33 TCS fractions and the positive control was 400 ppm metformin. Both diluted in DMEM with 0.5 % final DMSO concentration, incubated for 24 hours at 37 °C incubator in a humidified 5% CO2 atmosphere. The normal control was incubated with the solvent only. The cells were then incubated by acute high glucose-high insulin incubation for 5 minutes.

#### 2.3.1. pIRS1 ser312 measurement

The serine-312 phosphorylated IRS1 (pIRS1 ser312) was measured using in cell ELISA assay following manufacture’s manual guide (MBS9501484). Briefly, cells were fixated in plate base, following overnight incubation with primary antibodies, incubation using HRP conjugated secondary antibodies, addition of substrate, and reading the optical density (OD) value at 450 nm. The measured 450 OD was normalized using OD value of crystal violet staining that indicated the cell density in each well. The inhibition of ser-312 pIRS1 was measured relative to negative control myotube cells.

#### 2.3.2. Translocated GLUT4 measurement

Insulin resistant induced cells in round cover slip were treated as described above followed by fixation in 4% paraformaldehydes for 15 minutes. GLUT4 conjugated Alexa fluor 488 antibody (Rat IgG1, Santa Cruz SC-53566 AF488) was applied to the cells at a dilution of 1:50 in 5% FBS overnight in 4 °C. DAPI (4’,6-diamidino-2-phenylindole) staining for cell nuclei was conducted by 10 minutes incubating at 20 μg/mL after washing with PBS. The coverslips than washed in PBS and mounted before confocal imaging (Bradley et al., 2014). Image quantifications were conducted for each slide with two slides replication for each group. At least 5 images were captured per slide, therefore for each group at least 10 images were analyzed. For quantification of GLUT4 fluorescence intensity was carried out using Fiji ImageJ and was kept consistent between images. GLUT4 mean fluorescence intensity (MFI) was quantified by measuring the signal intensity after the background reduction for each image. No image processing was carried out prior to intensity analysis.

#### 2.3.3. Glycogen content measurement

Glycogen content was measured using glycogen ELISA kit following manufacture’s manual guide (Mybiosource, MBS729293). Briefly cells were harvested and lysed by three times freeze-thaw cycle. The supernatant then subjected to well coated primary antibody, adding the conjugate, and substrates respectively. The glycogen content was measured based on OD value of the standard curve.

### 2.4. Untargeted metabolomics analysis

### 2.4.1. Instrumentation and software

We used Q-Exactive hybrid quadrupole-orbitrap mass spectrometer (Thermo Fisher Scientific, Waltham, MA USA) to separate the metabolites of TCS. The metabolite quantification analysis was conducted using Thermo XCalibur version 4.2.28.14 and Compound Discoverer 3.3 (Thermo Fisher Scientific, Waltham, MA USA). The multivariate analysis was performed using SIMCA 16 (Sartorius, Goettingen, Germany).

### 2.4.2. Metabolites identification

Freeze dried TCS fractions (n=33) were dissolved in 1 mL of methanol using an ultrasonic water bath at room temperature for 10 min. The mixtures underwent filtration using a 0.2 micrometer syringe filter and placed in autosampler vial. Subsequently, an injection of 5 µL from each mixture was introduced into a Q-Exactive hybrid quadrupole-orbitrap mass spectrometer.

We employed an Accucore C18 (100 × 2.1 mm, 1.5 µm) as the LC separation column, while the UV detector was set at 254 nm wavelength. Heated electrospray ionization was used as the ionization source and the Q-Orbitrap as mass analyzer. Collision energy levels of 18, 35, and 53 eV were used for fragmentation. Other experiments were as follows, spray voltage of 3.8 kV, 320 °C capillary temperature, sheath gas flow rate 15 mL/min, and auxiliary gas flow rate 3 mL/min. The delivery system flow rate was adjusted at 0.3 mL/min, the autosampler temperature was maintained at 10 °C, and the sample injection volume was 5 µL. The mobile phase consisted of 0.1% formic acid in water (A) and acetonitrile (A) as the gradient elution system. The total scan time was 32 min with 2 min of calibration. The scan was set for full MS/dd MS2 scan ranges from 100 to 1500 m/z in positive and negative mode.

The mass spectrometer data were processed and quantitively analyzed using Thermo XCalibur and Compound Discoverer 3.3 software. The minimum peak detection was set at 10,000. The steps are including selecting and aligning the mass spectra, detecting known compound using databases, detecting the unknown compounds, identify the unknown compound by fragmentation patterns of mass spectrum (MS and MS/MS), spectral matching using previous studies related to *Menispermaceae* family and predictive fragmentation algorithms using CFM-ID (https://cfmid.wishartlab.com/predict) (Wang et al., 2022). The databases used in this study were mzCloud, ChemSpider (Pence & Williams, 2010) and PubChem (Kim et al., 2019). The mass error value (10 ppm) was set as the filtering criteria.

### 2.5. Molecular docking

The ten main active metabolites based on metabolomics and multivariate data analysis (MVDA) would be studied using molecular docking. Its two dimensional structures were acquired from PubChem (https://pubchem.ncbi.nlm.nih.gov) (Kim et al., 2019). The crystal structure of GLUT4 as target proteins were downloaded from Research Collaboratory for Structural Bioinformatics (RCSB) (https://www.rcsb.org/) (PDB ID: 3PCU) (Berman, 2000; Y. Zhang et al., 2012). The protein’s 3 dimensional structures obtained from primal protein complex than removed its water content and separating the native ligand using PyMOL software. Subsequently, the pdbqt format files were transformed from PDB format files by AutoDock Tools. The docking area was set at 40×40×40 at the docking site following its native ligand position. The affinities of each compound to its target protein were simulated by molecular docking using AutoDockTools software (Morris et al., 2009). The docking validation was conducted by redocking its native ligand with RMSD value < 2 Amstrong from its reference crystal structure as the threshold (Astiani et al., 2023).

### 2.6. Statistical analysis

Quantitative data were presented as mean ± SD (standard deviation). Normality and homogeneity data were required before the ANOVA test. One way ANOVA tests were performed for each variable measurement. If there were significant difference between groups, data than analyzed using Tukey’s comparison test with multiple comparison to negative control group. Statistical tests were performed using GraphPad Prism (GraphPad Software Inc., San Diego, CA, USA). Correlation analysis between the three measured variables was also performed using GraphPad Prism through Pearson correlation analysis.

The multivariate analysis was conducted in unsupervised manner using principal component analysis (PCA) and orthogonal projection to latent square (OPLS) performed using SIMCA software 16.0 (Sartorius, Goettingen, Germany). OPLS analysis used to reduce the dimensionality of data sets that contain both predictor and response variables, while preserving as much of the variation in the data as possible (Triba et al., 2015). In this study, we use OPLS model to find a linear relationship between two matrices: a predictor matrix of mass spectrometric data from TCS fractions (X matrix) and a response matrix of inhibition of pIRS1 ser312, enhancing translocated GLUT4 and glycogen (Y matrix). Each of the measured variables was developed into its own OPLS model.

## 3. Result and discussion

### 3.1. *In vitro* effect of TCS on insulin resistant induced L6.C11 skeletal muscle cell

To examine the bioactivity of TCS fractions, we carried out an *in vitro* study to identify the most potent fraction as insulin sensitizer. We firstly developed a cell model of insulin resistance based on previous procedure (Huang et al., 2002). We chose L6.C11 skeletal muscle cells because they have sufficient expression of IRS1 and GLUT4, which can be well measured using the chosen method. It has been previously reported that hyperinsulinemia conditions lead to the development of insulin resistance in cell culture (Huang et al., 2002; Pryor et al., 2000). Chronic insulin treatment in cell culture has been reported to reduce 60-70% of insulin receptor (IR) tyrosine phosphorylation, phosphoinositide 3-kinase activity (PI3K), and Akt activity. *T. crispa* has been reported to have a multiple targets-multiple components mechanism for its insulin-sensitizing activity, with the prediction of main pathway being the PI3K/Akt pathway (Zuhri et al., 2022). The measurement of phosphorylated insulin receptor substrate 1 at serine312 (IRS1 ser312) inhibition, glycogen content, and translocated GLUT4 are considered to be respective variables for the insulin-sensitizing activity of TCS.

#### 3.1.1. Inhibitory activity of TCS on ser312 IRS1 phosphorylation

The alterations in pIRS1 ser312 in L6.C11 skeletal muscle cells were measured in the TCS treated groups using the in-cell ELISA assay. The phosphorylation of IRS1 in ser312 may induce several defects on insulin signaling, including dissociation of IRS1 from the receptor concomitant with IRS1 degradation and interrupts IRS1 binding to its downstream effectors (Boura-Halfon & Zick, 2009). Inhibition of serine phosphorylation was one of emerging indicator of insulin sensitization regarding its in-line effect of increasing AKT phosphorylation. Under conditions of low phosphorylation of ser312, insulin signaling will be activated, whereas under conditions of its high phosphorylation, insulin signaling will be inhibited (Copps & White, 2012). The ser-312 pIRS inhibitory properties of the TCS fractions differed in each fraction (Figure 1A and Supplementary material 1).

**Figure 1.**
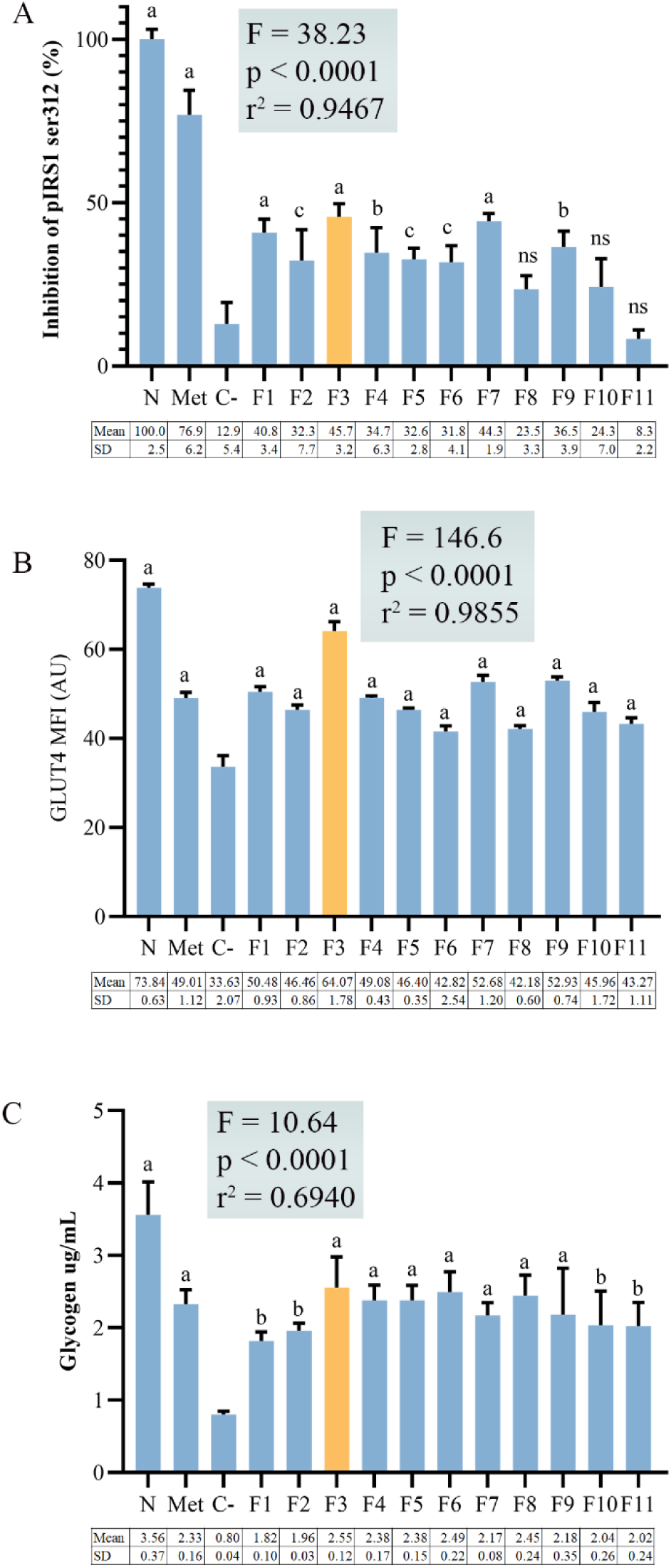
In vitro results of three measured variables. (A) pIRS1 ser312 inhibition, (B) translocated GLUT4, and (C) glycogen content. Treatment groups are denoted as F1 to F11, normal group as N, and metformin as Met. The treatment groups were compared to the negative control group (C-). a indicates significant difference with p-value < 0.0001; b indicates significant difference with p-value of 0.0001-0.0007; c indicates significant difference with p value 0.0023-0.0038; and ns indicates not significant (p>0.005). Statistics were performed using the one-way ANOVA test with a 95% confidence interval. The F, P, and r² values for each graph are shown at the top of the graph. Fraction F3 had the highest bioactivity of all three measured variables.

The one way ANOVA test showed a significant differences (p<0.0001; F=53.02) in inhibition activity of TCS fraction compared to the negative control (C-). The test showed that all treatment groups had a significant difference from the negative control except for treatment F8, F10, and F11 (p=0.2129; 0.1556; 0.9535). The C-group is a group of cells that were induced to insulin resistance and then incubated with only solvent. This group represents a high state of ser312 phosphorylation on IRS1, so a significant difference from this group is the expected treatment result. The post hoc Tukey’s Test result showed that the F3 was comparatively the most effective in reducing ser312 IRS1 phosphorylation than the other fractions. It reached 43.68 ± 3.20% inhibition, followed by fraction F7 (44.30 ± 1.92%) and fraction F1 (40.84 ± 4.40%). Fraction F3 from TCS is a fraction obtained from descending CPC fractionation at the early elution stage, which provides a gradient of polarity from polar to non-polar at the end of fractionation. Therefore, fraction F3 is a polar fraction. Based on the trend of bioactivity from the tested fractions, polar fractions (F1 to F7) tend to have higher activity than non-polar fractions. The active compounds responsible for the activity will be identified through OPLS multivariate analysis in section 3.3.

#### 3.1.2. Induce GLUT4 translocation to the cell membrane

The translocation of GLUT4 to cell membrane remains the key effect of insulin signaling that deployed to glucose uptake to the intracellular environment. The embedded GLUT4 in cell membrane is a requirement for glucose uptake from the circulation. Immunocytochemistry technique using GLUT4 specific antibody conjugated with Alexa fluor 488 (AF488) was conducted to reveal the increasing amount of translocated GLUT4 in cell membrane. The immunofluorescence was observed under confocal microscope at the same fluorescence wavelength using 63x magnifying factor.

The one-way ANOVA test results revealed significant differences among the treatment groups, as presented in Figure 1B and supplementary material 2. Significant differences (p< 0.05) in translocated GLUT4 compared to negative control were found in every TCS fraction with differ potency. In the previous study, it has been reported that GLUT4 transcription was unaltered when the glucose uptake was elevated in insulin resistant L6.C11 cells treated with *T. crispa*, measured by RT-PCR method (Noipha & Ninlaaesong, 2011).

Our findings showed a significant increase of GLUT4 that embedded to cell membrane as a result of TCS treatment. Further research is needed to investigate whether the increased translocation of GLUT4 is influenced by the level of GLUT4 transcription. The mean fluorescence intensity effected by F3 incubation was the most effective in inducing GLUT4 translocation to cell membrane (64.07 ± 1.78 AU) compared to the other fraction and even higher than the metformin treatment (49.01 ± 1.12 AU) at the same concentration (400 ppm). The representative images of confocal observation are shown in Figure 2 and the complete images were shown at Supplementary material 4.

**Figure 2.**
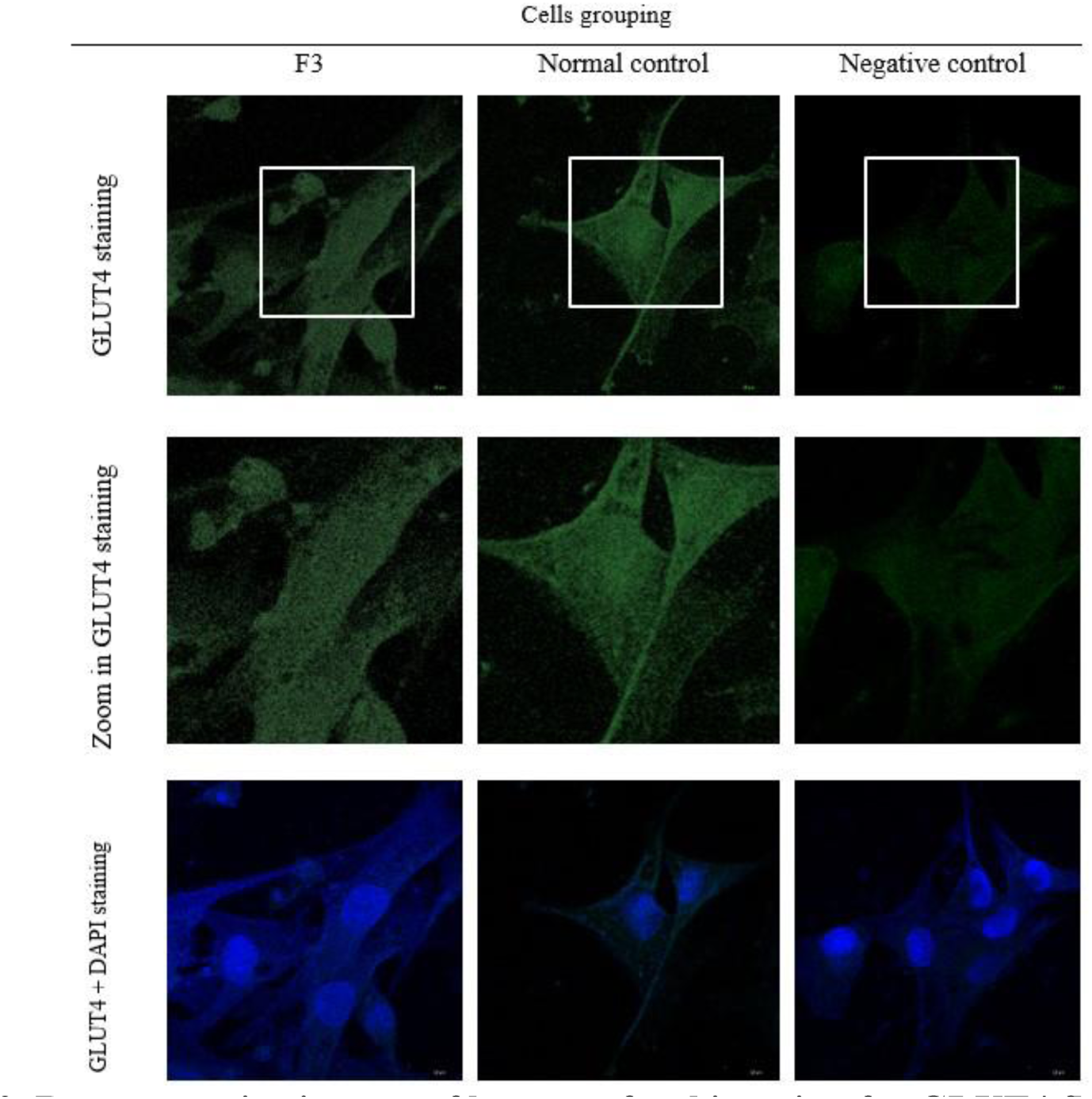
Representative images of laser confocal imaging for GLUT4 fluorescence. Quantification was conducted at 63x magnification from three experimental groups. The three images shown represent the results of fluorescence observations of GLUT4, with the highest value (normal control), lowest (negative control), and one active fraction (F3). The F3 group represents insulin-resistant cells treated with F3 at 400 ppm, while the negative group remains untreated. The normal group consists of uninduced cells treated only with the solvent used in F3 (0.5% (v/v) DMSO in DMEM. Images were captured at 490 nm for GLUT4 imaging and 510 nm for DAPI imaging. The displayed images have been adjusted for contrast to enhance fluorescence visibility. However, for fluorescence quantification using ImageJ, the original unaltered images were used.

#### 3.1.2. Activity of TCS as glycogen enhancer

The outcome effect of insulin signaling in cell culture was measured using the glycogen production in the L6.C11 skeletal muscle cells. All the TCS fractions showed significant induction of glycogen production compared to the negative control (Figure 1C and supplementary material 3). Our result showed that the F3 (2.55 ± 0.12 μg/mL) was the most effective fraction to induce glycogen production compared to the other fractions. Its activity was even higher than the metformin treatment (2.33 ± 0.16 μg/mL) at the same concentration. Glycogen was the outcome product of insulin signaling to keep the glucose homeostasis. It was reported that glycogen concentration in the muscle was 1-2% of the muscle mass approximately 400 g in a 70 kg person (Knuiman et al., 2015). A limitation of this study is that the biomass of L6.C11 cell culture was not measured, so no data was available to reflect muscle mass. The glycogen content of 2.55 ± 0.12 μg/mL in the F3 treatment showed a three-fold increase compared to the negative control, indicating the positive effect of TCS in attenuating insulin resistance conditions.

#### 3.1.3. Correlation of inhibition of pIRS1 ser312, translocated GLUT4, and glycogen content

Pearson correlation analysis was performed using GraphPad Prism software to investigate the relationship between the three variables. Normality and homogeneity tests were conducted prior to correlation analysis. The results of the normality test using Shapiro-Wilk test showed that the data was normally distributed (p>0.05) and Levene’s homogeneity of variances test showed that the data was homogeneous with a p-value>0.05. The results of the Pearson correlation test showed a positive and significant correlation between pIRS1 ser312 inhibition and GLUT4 translocation (p<0.05) with 59% coefficient determination value (Figure 3).

**Figure 3.**
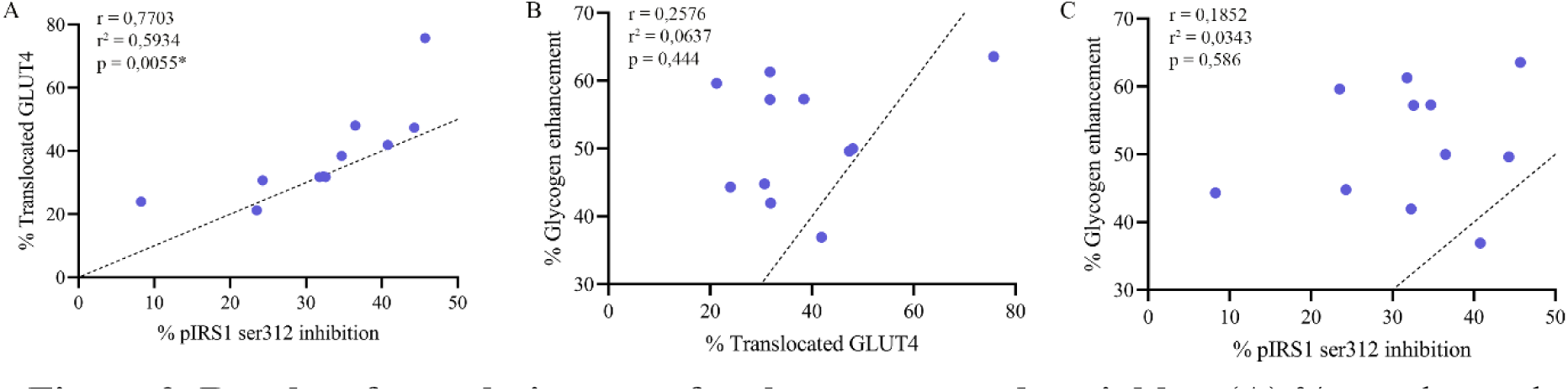
Results of correlation tests for three measured variables. (A) % translocated GLUT4 versus % pIRS1 ser312 inhibition, (B) % glycogen enhancement versus % translocated GLUT4, and (C) % glycogen enhancement versus % pIRS1 ser312 inhibition. The dotted line indicates the identity line, and the blue dots indicate the correlation between the two measured variables (X-axis and Y-axis). Statistics were performed using the Pearson correlation test with a 95% confidence interval. The r, r², and p values for each graph are shown at the top of the graph. A p-value with an asterisk indicates a significant correlation. A positive correlation was found between all three measured variables. A significant correlation was found between % pIRS1 ser312 inhibition and % translocated GLUT4, but neither variable had a significant correlation with % glycogen enhancement.

Conversely, the correlation between pIRS1 ser312 and glycogen, and translocated GLUT4 and glycogen, showed a positive but non-significant relationship. This suggests that other variables play a major role in regulating glycogen content such as the activity of glycogenesis and glycogen synthase enzymes involved in glycogen synthesis. Inhibition of pIRS1 ser312 contributes to the induction of GLUT4 vesicles to move to the cell membrane. GLUT4 then facilitates glucose entry into the intracellular environment. Glucose that has entered the cell is then converted into glycogen through a process initiated by the glycogenesis and glycogen synthase enzymes (Gual et al., 2005). The process is a complex signaling cascade, and inhibition of pIRS1 ser312 and GLUT4 translocation only has a small correlational contribution to glycogen content.

### 3.2. Putative identification of TCS metabolites

Putative identification of phytoconstituents present in TCS was performed in this study as explained in 2.4.2 subsection. Phytochemicals found in natural product were in a complex mixture at wide polarity differences (Q. W. Zhang et al., 2018). The CPC fractionation process was expected to produce slightly different compound combinations for each obtained fraction, which might have varied biological potency. A total of 2667 features were detected in both positive and negative ionization modes. By analyzing the MS1 and MS1 mass spectra of the analyte peaks, 58 metabolites were successfully identified using data from previous studies and databases (Table 2) (Ahmad et al., 2016; Bowen & Motawe, 1985; Choudhary, Ismail, Ali, et al., 2010; Choudhary, Ismail, Shaari, et al., 2010; Dong et al., 2010; Fukuda et al., 1985, 1993; Gao et al., 2016; M. Ismail & Choudhary, 2016; Kim et al., 2019; Lam et al., 2018, 2012; Li et al., 2004a, 2004b; Pachaly et al., 1992; Thomas et al., 2016; Van Kiem et al., 2010; Xu et al., 2017; Yonemitsu et al., 1995; Y. Zhang et al., 2010).

**Table 2.**
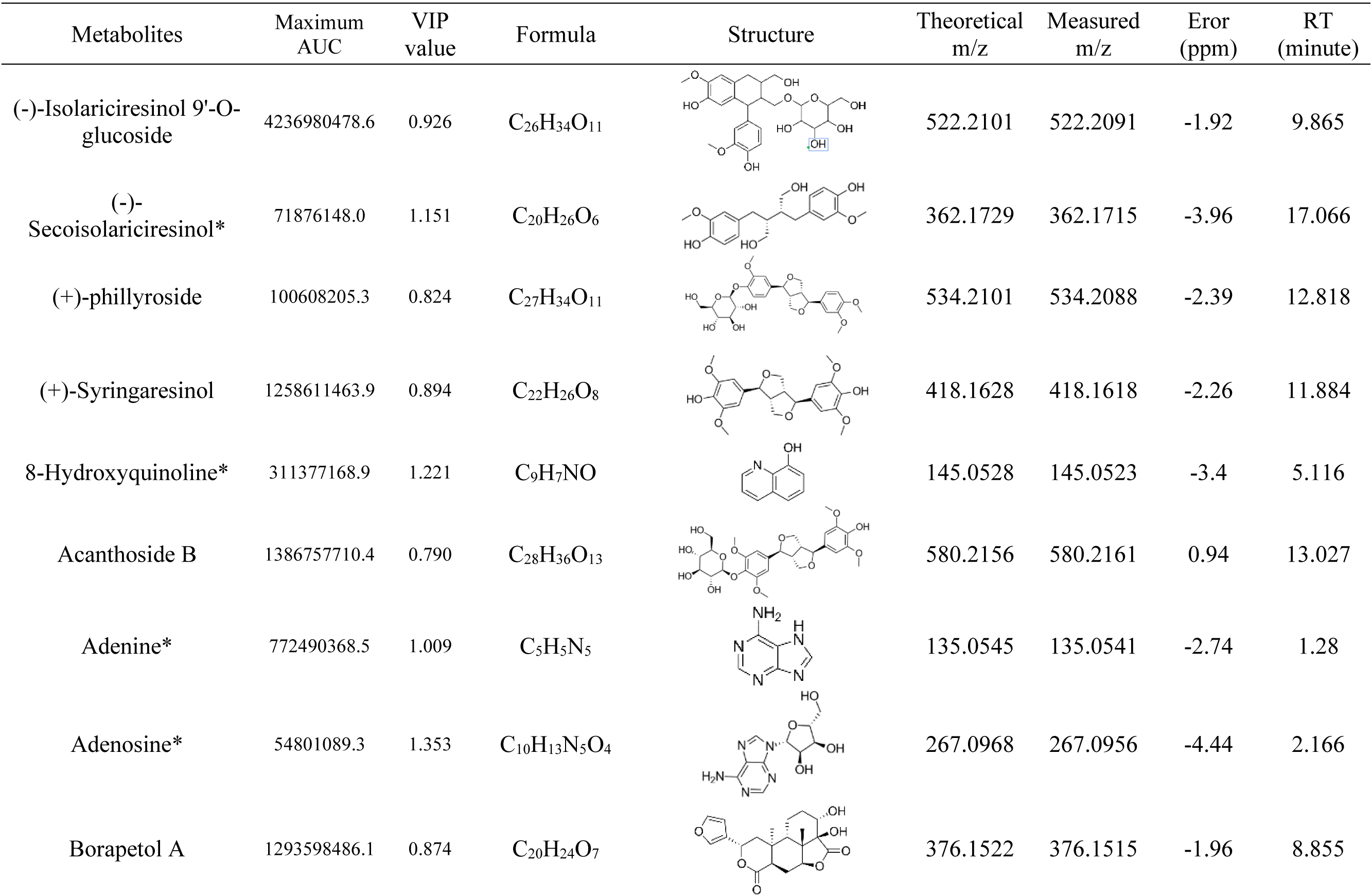
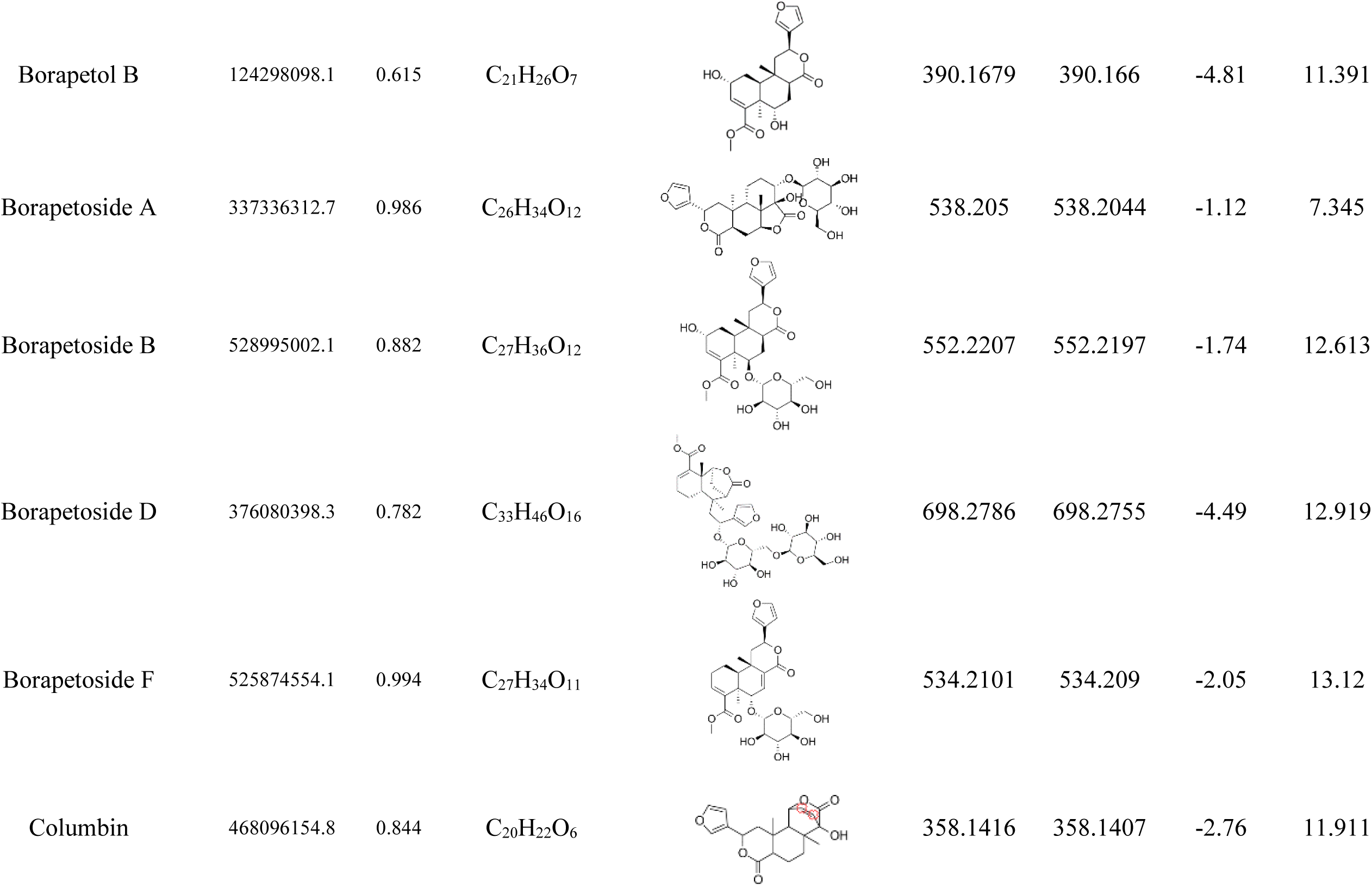
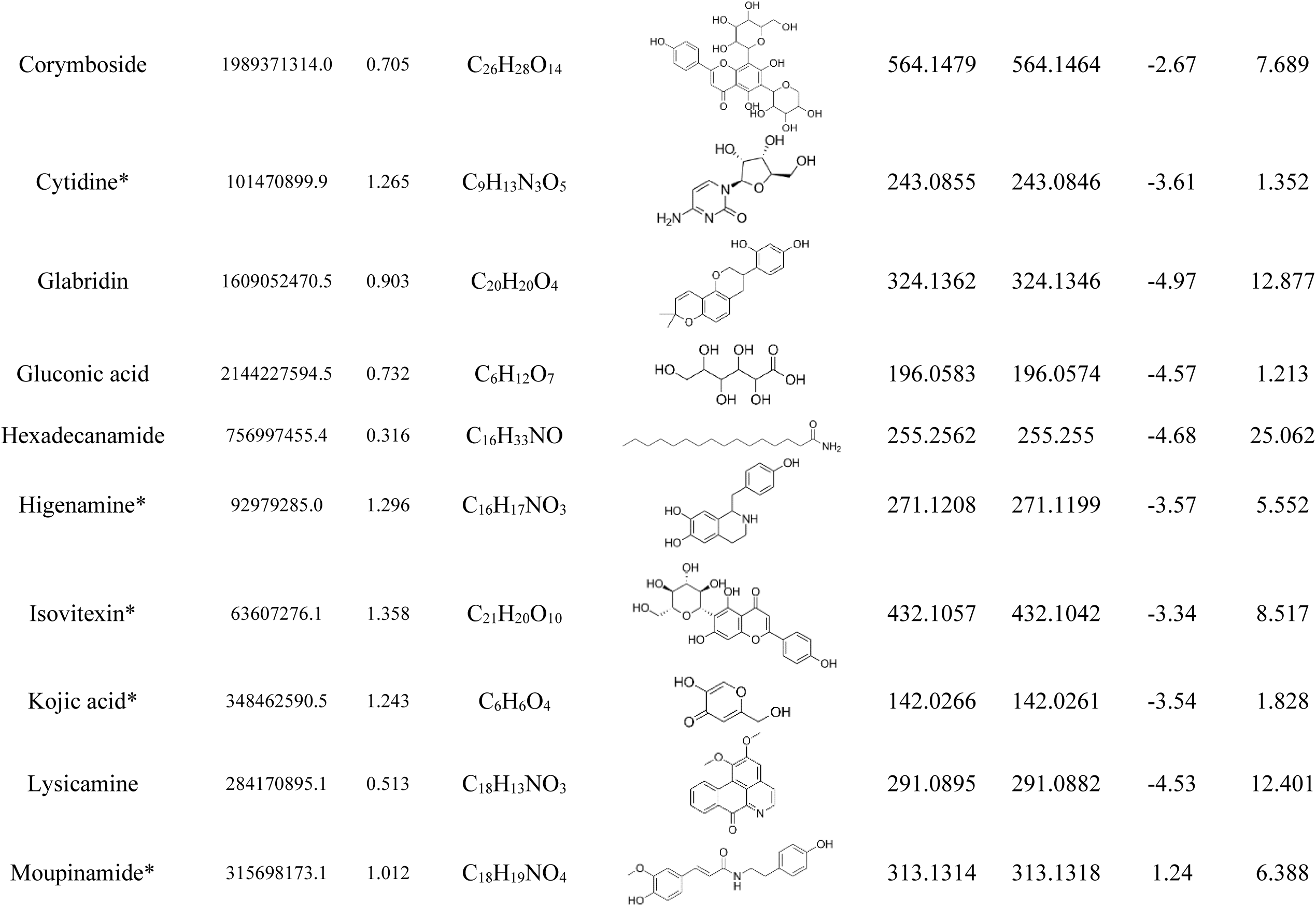
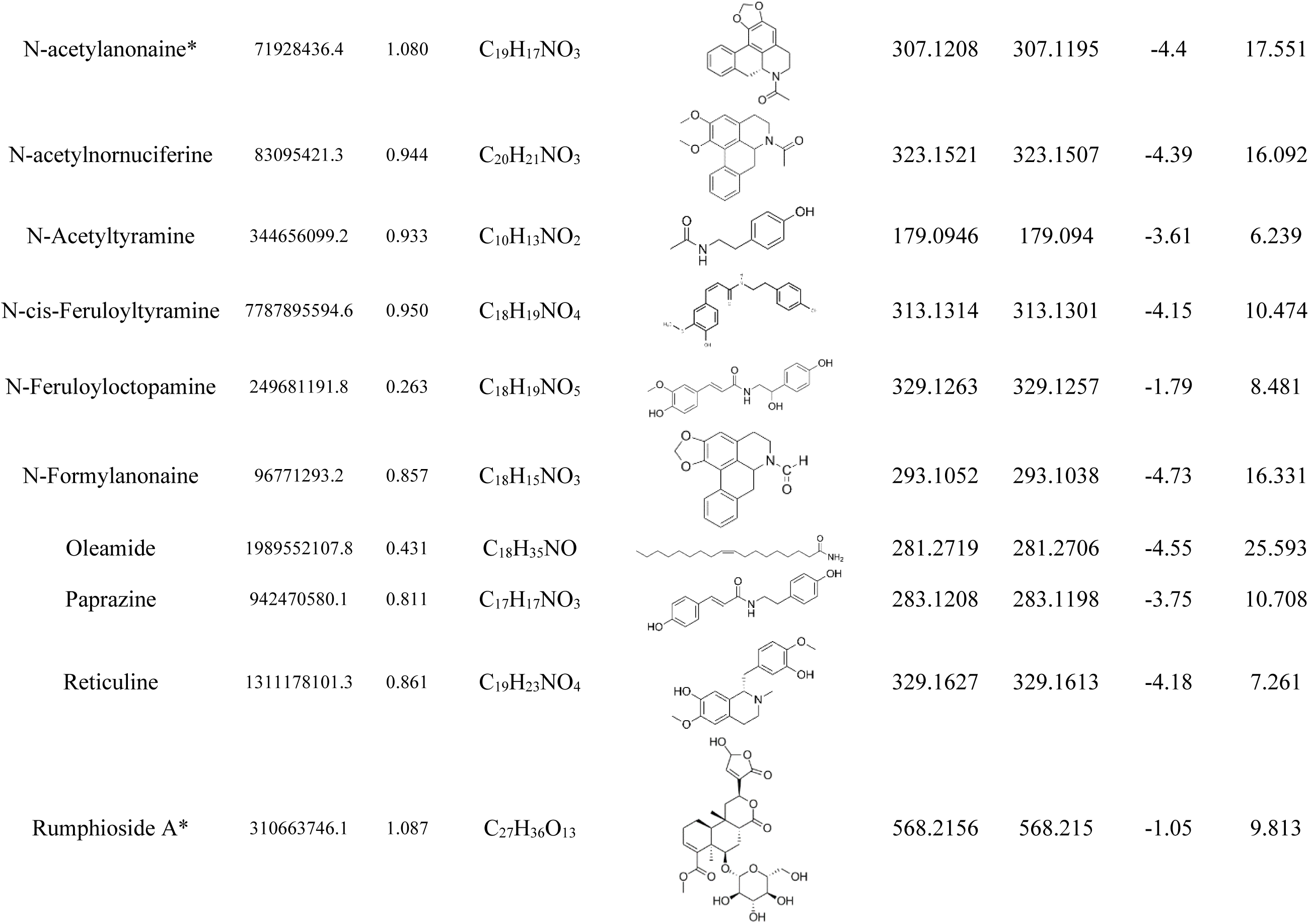
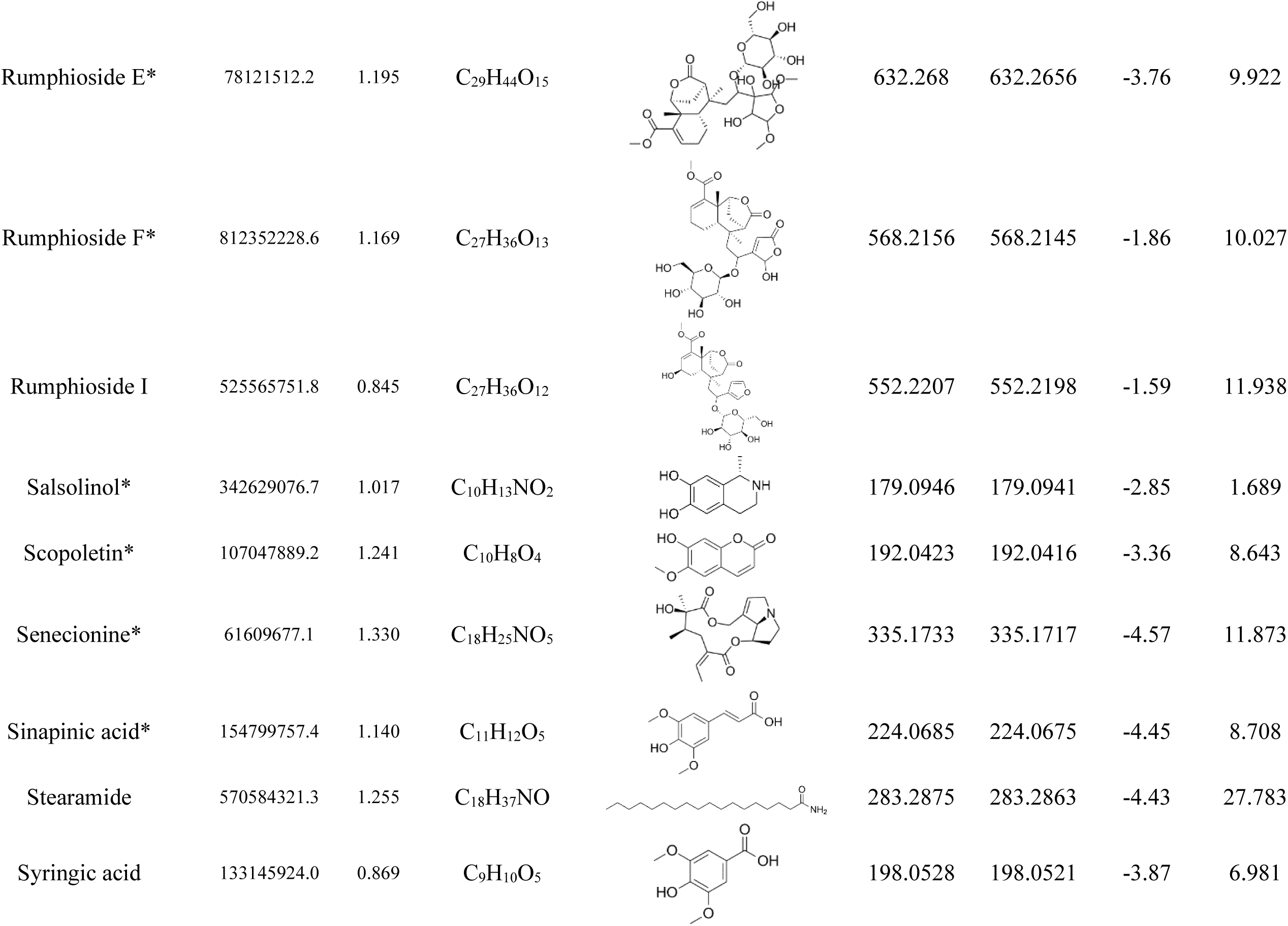
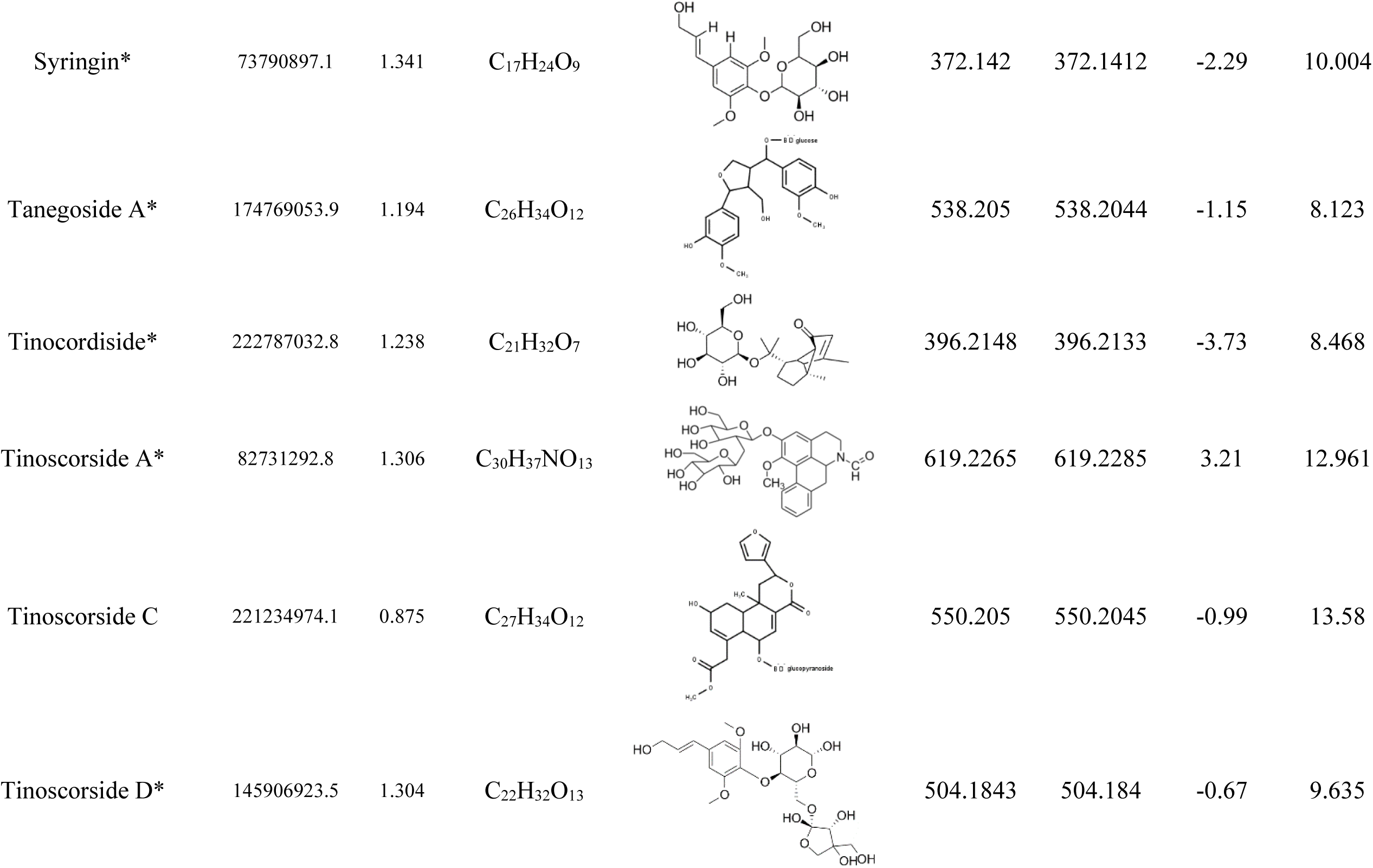
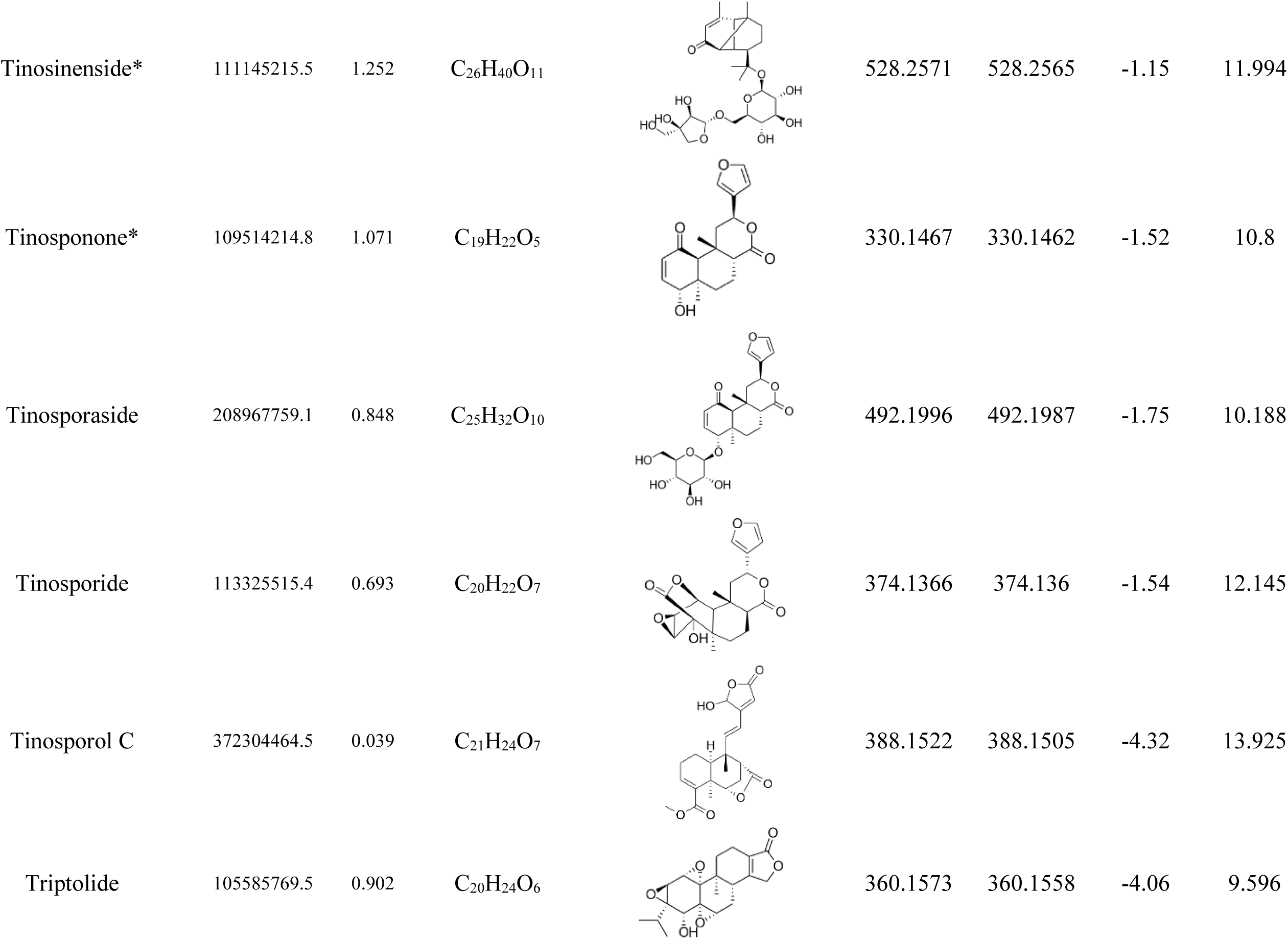
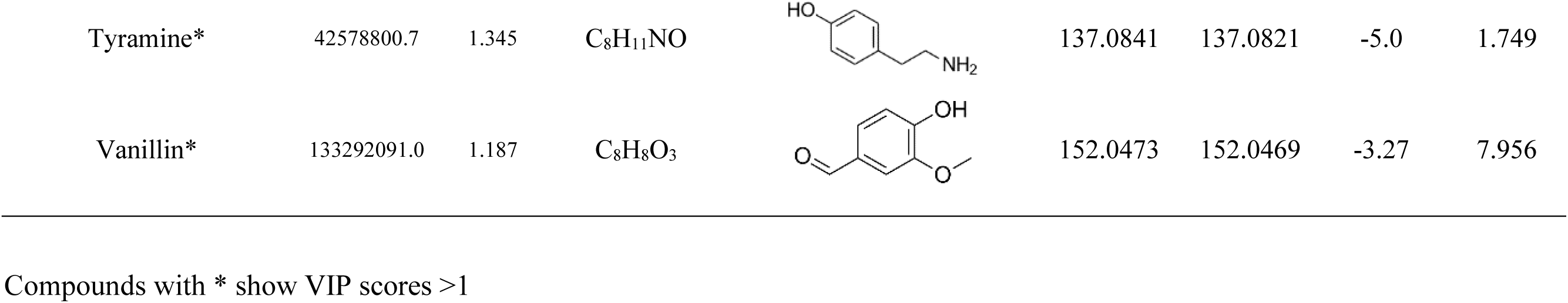
Identified metabolites of TCS.

| Metabolites | Maximum AUC | VIP value | Formula | Structure | Theoretical m/z | Measured m/z | Error (ppm) | RT (minute) |
| --- | --- | --- | --- | --- | --- | --- | --- | --- |
| (-)-Isolariciresinol 9'-O-glucoside | 4236980478.6 | 0.926 | C<sub>26</sub>H<sub>34</sub>O<sub>11</sub> |  | 522.2101 | 522.2091 | -1.92 | 9.865 |
| (-)-Secoisolariciresinol* | 71876148.0 | 1.151 | C<sub>20</sub>H<sub>26</sub>O<sub>6</sub> |  | 362.1729 | 362.1715 | -3.96 | 17.066 |
| (+)-phillyroside | 100608205.3 | 0.824 | C<sub>27</sub>H<sub>34</sub>O<sub>11</sub> |  | 534.2101 | 534.2088 | -2.39 | 12.818 |
| (+)-Syringaresinol | 1258611463.9 | 0.894 | C<sub>22</sub>H<sub>26</sub>O<sub>8</sub> |  | 418.1628 | 418.1618 | -2.26 | 11.884 |
| 8-Hydroxyquinoline* | 311377168.9 | 1.221 | C<sub>9</sub>H<sub>7</sub>NO |  | 145.0528 | 145.0523 | -3.4 | 5.116 |
| Acanthoside B | 1386757710.4 | 0.790 | C<sub>28</sub>H<sub>36</sub>O<sub>13</sub> |  | 580.2156 | 580.2161 | 0.94 | 13.027 |
| Adenine* | 772490368.5 | 1.009 | C<sub>5</sub>H<sub>5</sub>N<sub>5</sub> |  | 135.0545 | 135.0541 | -2.74 | 1.28 |
| Adenosine* | 54801089.3 | 1.353 | C<sub>10</sub>H<sub>13</sub>N<sub>5</sub>O<sub>4</sub> |  | 267.0968 | 267.0956 | -4.44 | 2.166 |
| Borapetol A | 1293598486.1 | 0.874 | C<sub>20</sub>H<sub>24</sub>O<sub>7</sub> |  | 376.1522 | 376.1515 | -1.96 | 8.855 |
| Borapetol B | 124298098.1 | 0.615 | C<sub>21</sub>H<sub>26</sub>O<sub>7</sub> |  | 390.1679 | 390.166 | -4.81 | 11.391 |
| Borapetoside A | 337336312.7 | 0.986 | C<sub>26</sub>H<sub>34</sub>O<sub>12</sub> |  | 538.205 | 538.2044 | -1.12 | 7.345 |
| Borapetoside B | 528995002.1 | 0.882 | C<sub>27</sub>H<sub>36</sub>O<sub>12</sub> |  | 552.2207 | 552.2197 | -1.74 | 12.613 |
| Borapetoside D | 376080398.3 | 0.782 | C<sub>33</sub>H<sub>46</sub>O<sub>16</sub> |  | 698.2786 | 698.2755 | -4.49 | 12.919 |
| Borapetoside F | 525874554.1 | 0.994 | C<sub>27</sub>H<sub>34</sub>O<sub>11</sub> |  | 534.2101 | 534.209 | -2.05 | 13.12 |
| Columbin | 468096154.8 | 0.844 | C<sub>20</sub>H<sub>22</sub>O<sub>6</sub> |  | 358.1416 | 358.1407 | -2.76 | 11.911 |
| Corymboside | 1989371314.0 | 0.705 | C<sub>26</sub>H<sub>28</sub>O<sub>14</sub> |  | 564.1479 | 564.1464 | -2.67 | 7.689 |
| Cytidine* | 101470899.9 | 1.265 | C<sub>9</sub>H<sub>13</sub>N<sub>3</sub>O<sub>5</sub> |  | 243.0855 | 243.0846 | -3.61 | 1.352 |
| Glabridin | 1609052470.5 | 0.903 | C<sub>20</sub>H<sub>20</sub>O<sub>4</sub> |  | 324.1362 | 324.1346 | -4.97 | 12.877 |
| Gluconic acid | 2144227594.5 | 0.732 | C<sub>6</sub>H<sub>12</sub>O<sub>7</sub> |  | 196.0583 | 196.0574 | -4.57 | 1.213 |
| Hexadecanamide | 756997455.4 | 0.316 | C<sub>16</sub>H<sub>33</sub>NO |  | 255.2562 | 255.255 | -4.68 | 25.062 |
| Higenamine* | 92979285.0 | 1.296 | C<sub>16</sub>H<sub>17</sub>NO<sub>3</sub> |  | 271.1208 | 271.1199 | -3.57 | 5.552 |
| Isovitexin* | 63607276.1 | 1.358 | C<sub>21</sub>H<sub>20</sub>O<sub>10</sub> |  | 432.1057 | 432.1042 | -3.34 | 8.517 |
| Kojic acid* | 348462590.5 | 1.243 | C<sub>6</sub>H<sub>6</sub>O<sub>4</sub> |  | 142.0266 | 142.0261 | -3.54 | 1.828 |
| Lysicamine | 284170895.1 | 0.513 | C<sub>18</sub>H<sub>13</sub>NO<sub>3</sub> |  | 291.0895 | 291.0882 | -4.53 | 12.401 |
| Moupinamide* | 315698173.1 | 1.012 | C<sub>18</sub>H<sub>19</sub>NO<sub>4</sub> |  | 313.1314 | 313.1318 | 1.24 | 6.388 |
| N-acetylanonaine* | 71928436.4 | 1.080 | C<sub>19</sub>H<sub>17</sub>NO<sub>3</sub> |  | 307.1208 | 307.1195 | -4.4 | 17.551 |
| N-acetylnornuciferine | 83095421.3 | 0.944 | C<sub>20</sub>H<sub>21</sub>NO<sub>3</sub> |  | 323.1521 | 323.1507 | -4.39 | 16.092 |
| N-Acetyltyramine | 344656099.2 | 0.933 | C<sub>10</sub>H<sub>13</sub>NO<sub>2</sub> |  | 179.0946 | 179.094 | -3.61 | 6.239 |
| N-cis-Feruloyltyramine | 7787895594.6 | 0.950 | C<sub>18</sub>H<sub>19</sub>NO<sub>4</sub> |  | 313.1314 | 313.1301 | -4.15 | 10.474 |
| N-Feruloyloctopamine | 249681191.8 | 0.263 | C<sub>18</sub>H<sub>19</sub>NO<sub>5</sub> |  | 329.1263 | 329.1257 | -1.79 | 8.481 |
| N-Formylanonaine | 96771293.2 | 0.857 | C<sub>18</sub>H<sub>15</sub>NO<sub>3</sub> |  | 293.1052 | 293.1038 | -4.73 | 16.331 |
| Oleamide | 1989552107.8 | 0.431 | C<sub>18</sub>H<sub>35</sub>NO |  | 281.2719 | 281.2706 | -4.55 | 25.593 |
| Paprazine | 942470580.1 | 0.811 | C<sub>17</sub>H<sub>17</sub>NO<sub>3</sub> |  | 283.1208 | 283.1198 | -3.75 | 10.708 |
| Reticuline | 1311178101.3 | 0.861 | C<sub>19</sub>H<sub>23</sub>NO<sub>4</sub> |  | 329.1627 | 329.1613 | -4.18 | 7.261 |
| Rumphioside A* | 310663746.1 | 1.087 | C<sub>27</sub>H<sub>36</sub>O<sub>13</sub> |  | 568.2156 | 568.215 | -1.05 | 9.813 |
| Rumphioside E* | 78121512.2 | 1.195 | C<sub>29</sub>H<sub>44</sub>O<sub>15</sub> |  | 632.268 | 632.2656 | -3.76 | 9.922 |
| Rumphioside F* | 812352228.6 | 1.169 | C<sub>27</sub>H<sub>36</sub>O<sub>13</sub> |  | 568.2156 | 568.2145 | -1.86 | 10.027 |
| Rumphioside I | 525565751.8 | 0.845 | C<sub>27</sub>H<sub>36</sub>O<sub>12</sub> |  | 552.2207 | 552.2198 | -1.59 | 11.938 |
| Salsolinol* | 342629076.7 | 1.017 | C<sub>10</sub>H<sub>13</sub>NO<sub>2</sub> |  | 179.0946 | 179.0941 | -2.85 | 1.689 |
| Scopoletin* | 107047889.2 | 1.241 | C<sub>10</sub>H<sub>8</sub>O<sub>4</sub> |  | 192.0423 | 192.0416 | -3.36 | 8.643 |
| Senecionine* | 61609677.1 | 1.330 | C<sub>18</sub>H<sub>25</sub>NO<sub>5</sub> |  | 335.1733 | 335.1717 | -4.57 | 11.873 |
| Sinapinic acid* | 154799757.4 | 1.140 | C<sub>11</sub>H<sub>12</sub>O<sub>5</sub> |  | 224.0685 | 224.0675 | -4.45 | 8.708 |
| Stearamide | 570584321.3 | 1.255 | C<sub>18</sub>H<sub>37</sub>NO |  | 283.2875 | 283.2863 | -4.43 | 27.783 |
| Syringic acid | 133145924.0 | 0.869 | C<sub>9</sub>H<sub>10</sub>O<sub>5</sub> |  | 198.0528 | 198.0521 | -3.87 | 6.981 |
| Syringin* | 73790897.1 | 1.341 | C<sub>17</sub>H<sub>24</sub>O<sub>9</sub> |  | 372.142 | 372.1412 | -2.29 | 10.004 |
| Tanegoside A* | 174769053.9 | 1.194 | C<sub>26</sub>H<sub>34</sub>O<sub>12</sub> |  | 538.205 | 538.2044 | -1.15 | 8.123 |
| Tinocordiside* | 222787032.8 | 1.238 | C<sub>21</sub>H<sub>32</sub>O<sub>7</sub> |  | 396.2148 | 396.2133 | -3.73 | 8.468 |
| Tinoscorside A* | 82731292.8 | 1.306 | C<sub>30</sub>H<sub>37</sub>NO<sub>13</sub> |  | 619.2265 | 619.2285 | 3.21 | 12.961 |
| Tinoscorside C | 221234974.1 | 0.875 | C<sub>27</sub>H<sub>34</sub>O<sub>12</sub> |  | 550.205 | 550.2045 | -0.99 | 13.58 |
| Tinoscorside D* | 145906923.5 | 1.304 | C<sub>22</sub>H<sub>32</sub>O<sub>13</sub> |  | 504.1843 | 504.184 | -0.67 | 9.635 |
| Tinosinenside* | 111145215.5 | 1.252 | C<sub>26</sub>H<sub>40</sub>O<sub>11</sub> |  | 528.2571 | 528.2565 | -1.15 | 11.994 |
| Tinosponone* | 109514214.8 | 1.071 | C<sub>19</sub>H<sub>22</sub>O<sub>5</sub> |  | 330.1467 | 330.1462 | -1.52 | 10.8 |
| Tinosporaside | 208967759.1 | 0.848 | C<sub>25</sub>H<sub>32</sub>O<sub>10</sub> |  | 492.1996 | 492.1987 | -1.75 | 10.188 |
| Tinosporide | 113325515.4 | 0.693 | C<sub>20</sub>H<sub>22</sub>O<sub>7</sub> |  | 374.1366 | 374.136 | -1.54 | 12.145 |
| Tinosporol C | 372304464.5 | 0.039 | C<sub>21</sub>H<sub>24</sub>O<sub>7</sub> |  | 388.1522 | 388.1505 | -4.32 | 13.925 |
| Triptolide | 105585769.5 | 0.902 | C<sub>20</sub>H<sub>24</sub>O<sub>6</sub> |  | 360.1573 | 360.1558 | -4.06 | 9.596 |
| Tyramine* | 42578800.7 | 1.345 | C<sub>8</sub>H<sub>11</sub>NO |  | 137.0841 | 137.0821 | -5.0 | 1.749 |
| Vanillin* | 133292091.0 | 1.187 | C<sub>8</sub>H<sub>8</sub>O<sub>3</sub> |  | 152.0473 | 152.0469 | -3.27 | 7.956 |

Compounds with \* show VIP scores >1

Based on the PCA analysis (Figure 4), the TCS fractions were divided into four clusters regarding their chemical profile similarity. The first two main components remain presenting 88.2% of the total variance. It was showed that sequentially the group 1 to 5 had tinosporol C, n-feruloyloctopamine, n-feruloyloctopamine, borapetoside I, and columbin as their most abundant metabolites. Group 1 consists of F1 to F6 (A, B, and C), group 2 consists of F10 (A, B, and C) and F11 (A and C), group 3 consists of F7 and F9 (A, B, and C), group 4 consists of F8A, B, and C, and group 5 only consist of F11B. The grouped fractions indicate the similarity of their chemical profile. It is indicated that the fractions have different compositions.

**Figure 4.**
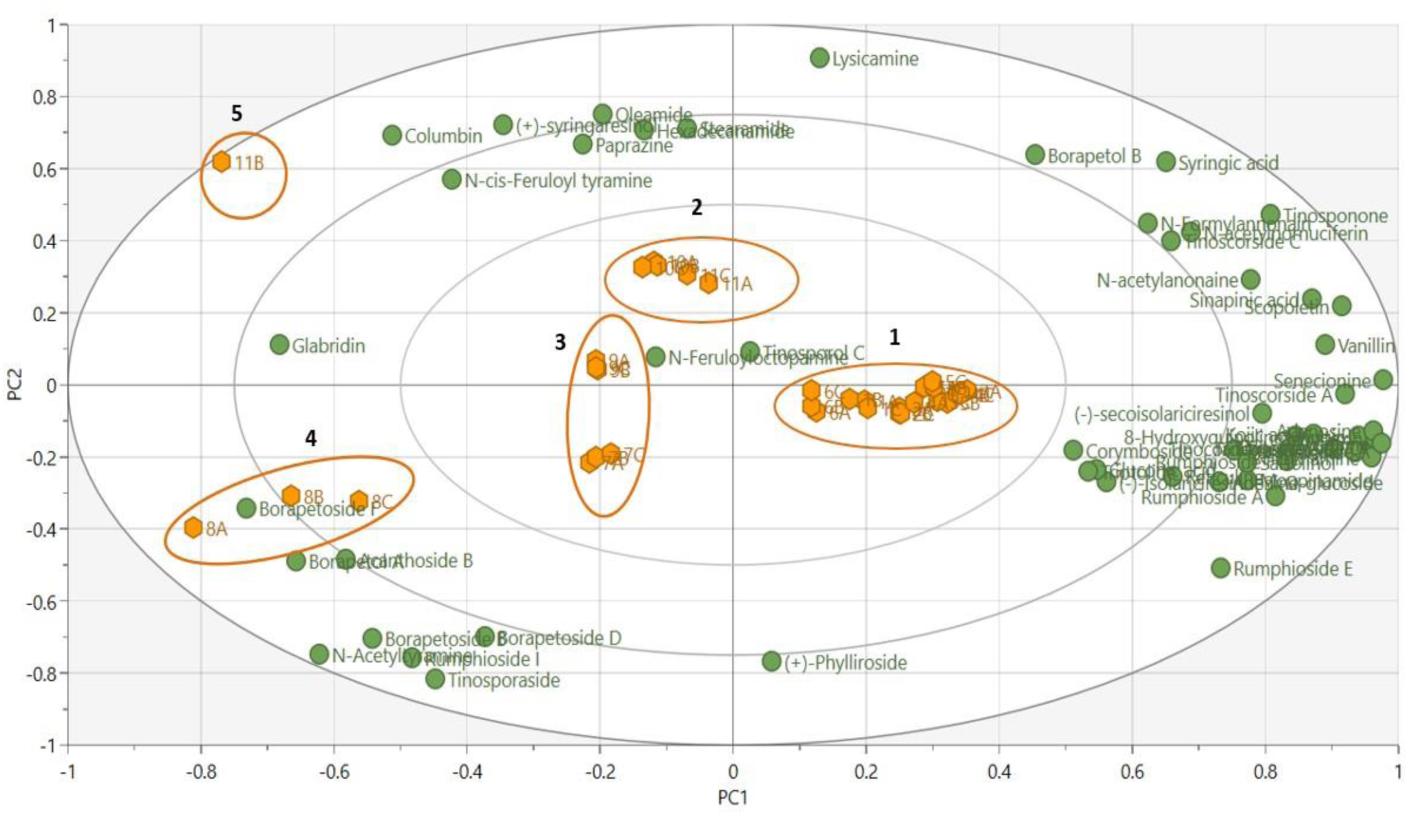
Scatter bi-plot of principal component analysis (PCA) of two principal components. The X-axis represents the projection of the data onto principal component 1 (PC1) and the Y-axis onto principal component 2 (PC2). Both indicate the proportion of the variance explained by the first and second predictive component. The green points represent the identified metabolites, the orange points represent the TCS fractions, and the ellipses represent the TCS fractions that are clustered based on the similarity of their metabolite profiles. The first two main components remain presenting 88.2% of the total variance. There are five clusters of TCS fractions based on the similarity of their metabolite profiles.

Phytochemicals, as other chemicals have a selective of solubility based on its polarity or stated as “like dissolves like”. Polar chemicals are more likely to solved in polar solvent and vice versa (Lefebvre et al., 2021; Smith et al., 1977). Our work was conducted CPC fractionation using H Arizona solvent system (n hexane-ethyl acetate-methanol-water; 1-3-1-3) those solvents created two immiscible biphasic solvents. The lower phase or aqueous phase remains more polar, and the upper phase or organic phase remains nonpolar. The solvent mixture saturates each other, constructing a unique solvent composition. The polar phase from Arizona system type H consist of 67.2 water, 11.9 ethyl acetate, 0.04 hexane, and 20.9 methanol (% v/v) (Berthod et al., 2005). This polar solvent mixture would likely dissolve polar phytoconstituents from TCS methanol extract.

The identified metabolites were grouped in glycosides, flavonoids, alkaloids, coumarins, and nucleotides groups. Therefore, it did not directly represent the most active metabolites as insulin sensitizer. To identify the highly active metabolites, we conducted OPLS analysis to reveal the highly correlated metabolites to each measured bioactivity.

### 3.3. Identification of TCS active metabolites

The OPLS model was further established to identify the active biomarker of TCS in each measured bioactivity. Before further examination of OPLS model, the quality of the OPLS models were assessed (Figure 5). In general, R^2^Y and Q^2^ provide an estimation of fitness and quality of the model prediction. To achieve a high predictive ability, the values of R^2^Y and Q^2^ should be close to one. The OPLS model generates values of 0.955 and 0.939 suggesting that the model has excellent reliability and predictability (Triba et al., 2015).

**Figure 5.**
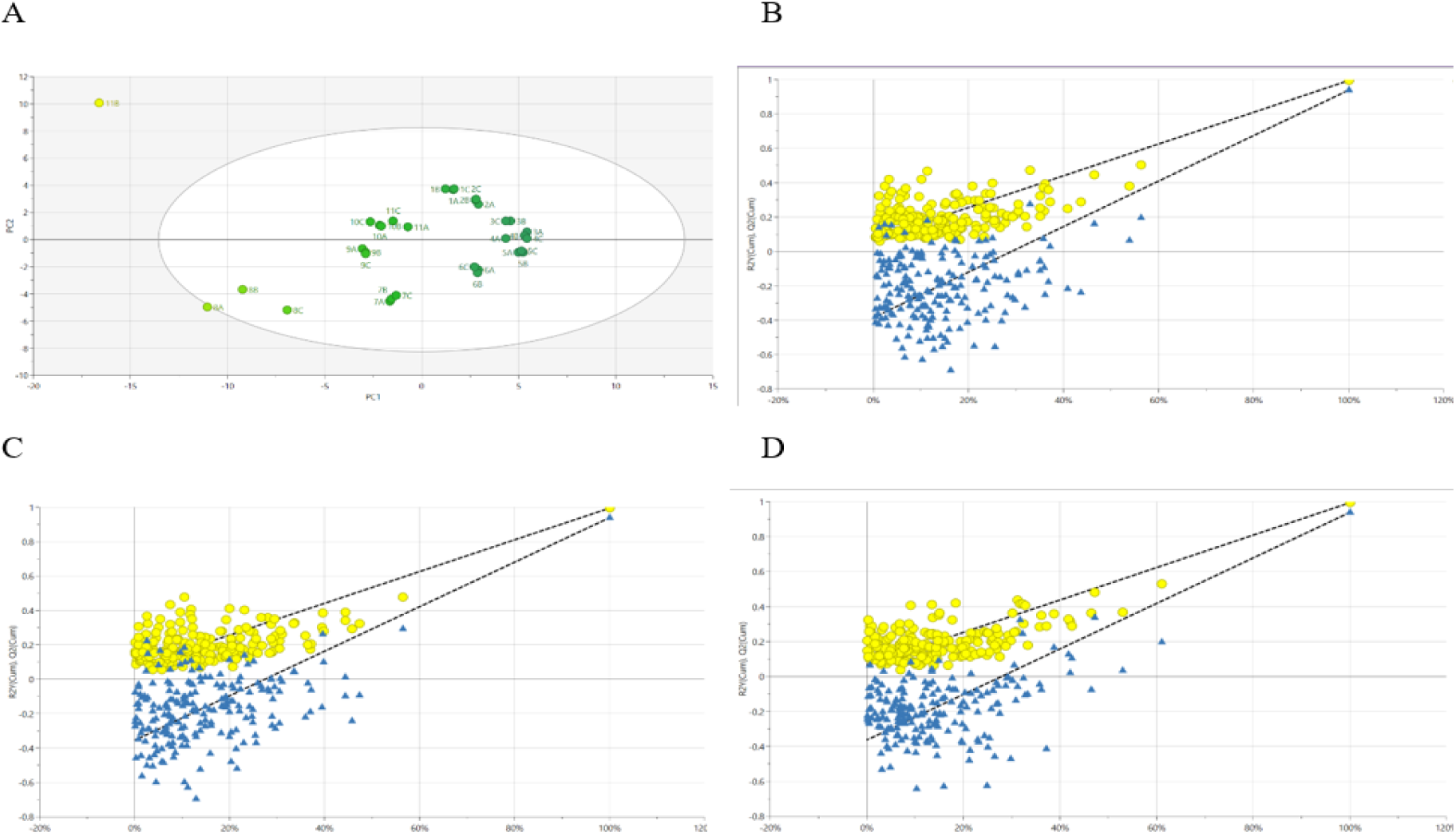
The quality control of the OPLS models. (**A**) Score scatter plot of the OPLS model for all TCS fractions (R^2^X= 0.636, R^2^Y= 0.955, Q^2^= 0.939). (**B-D**) Permutation test plot with 100 cycles. The yellow dots represent R^2^Y of model matrix information; the blue dots represent Q^2^ the predictive power of the original model. In consecutive order, the data represents inhibition of pIRS1 ser312, increased translocation of GLUT4, and increased glycogen levels.

The OPLS analysis results, shown as scatter plot (Figure 6), show that the TCS fractions have a clustering based on their insulin sensitizing activity against L6.C11 skeletal muscle cell culture. The first two principal components (PCs) explain 76.4% of the total variance in the OPLS model. The OPLS analysis results indicate that the fractions with the highest activity, in consecutive order, are fraction 3 (A, B, and C), fraction 1A, fraction 5 (B and C), and fraction 4B. It seems that the solvent polarity contributes to the separation of samples, which is related to their activity. The TCS polar fractions tend to be more active than the less polar fraction.

**Figure 6.**
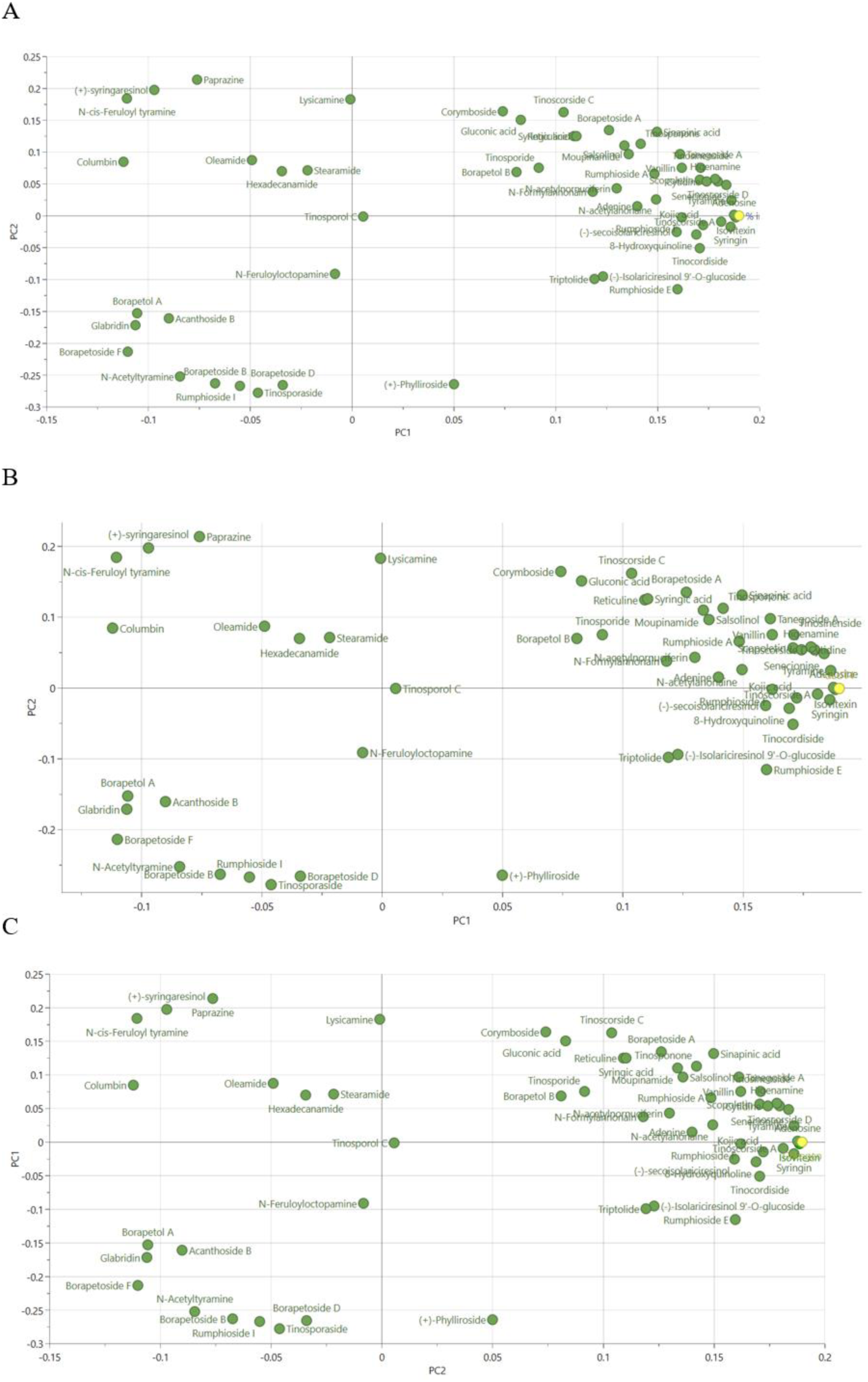
The OPLS loading plots. (A) inhibition of pIRS1 ser312, (B) translocated GLUT4, and (C) Glycogen content. The X-axis represents the projection of the data onto principal component 1 (PC1) and the Y-axis onto principal component 2 (PC2). Both indicate the proportion of the variance explained by the first and second predictive component and normalized in unit length. The green points represent the identified TCS metabolites, the yellow points represent the measured bioactivities. The position of the metabolite points to the right indicates a stronger correlation with bioactivity.

The results of the OPLS model were verified by a permutation test. At the three bioactivity dataset, shown that the Q^2^ point of the model from left to right is lower than the original Q^2^ point, and the R^2^ and Q^2^ values of the model are more than 0.9. Those which indicates that the model can reliably predict the results without any overfitting phenomenon (n=100) (Triba et al., 2015) (Figure 5).

To determine the metabolites with the highest contribution to bioactivity, the OPLS loading plot and Y-related profiles were under scrutinized. In the loading plot, the closer the coordinates of a metabolite to the zero coordinates (0,0), means the smaller of its contribution to the bioactivity (Figure 6). We also used Variable of Importance Projection (VIP) scores to estimate the contribution of each metabolite to the bioactivities as predicted by the OPLS model. The VIP scores > 1 was set as threshold for minimal contribution (Yuliana, Khatib, Verpoorte, & Choi, 2011). 26 metabolites reached this threshold as metabolites marked in Table 2. The Y-related coefficient plot indicated the correlation of metabolites to insulin sensitizer activity (Figure 7). The higher value regarded a higher correlation. Metabolites that showed the greatest contribution to bioactivity were isovitexin, syringin, tyramine, senesionine, tinoscorsida D, tinoscorsida A, higenamin, scopoletine, cytidine, and tinocordiside. The metabolites also pass the threshold for the VIP value. It can be concluded that all signals from the active compounds have a significant contribution to the insulin sensitizing activity.

**Figure 7.**
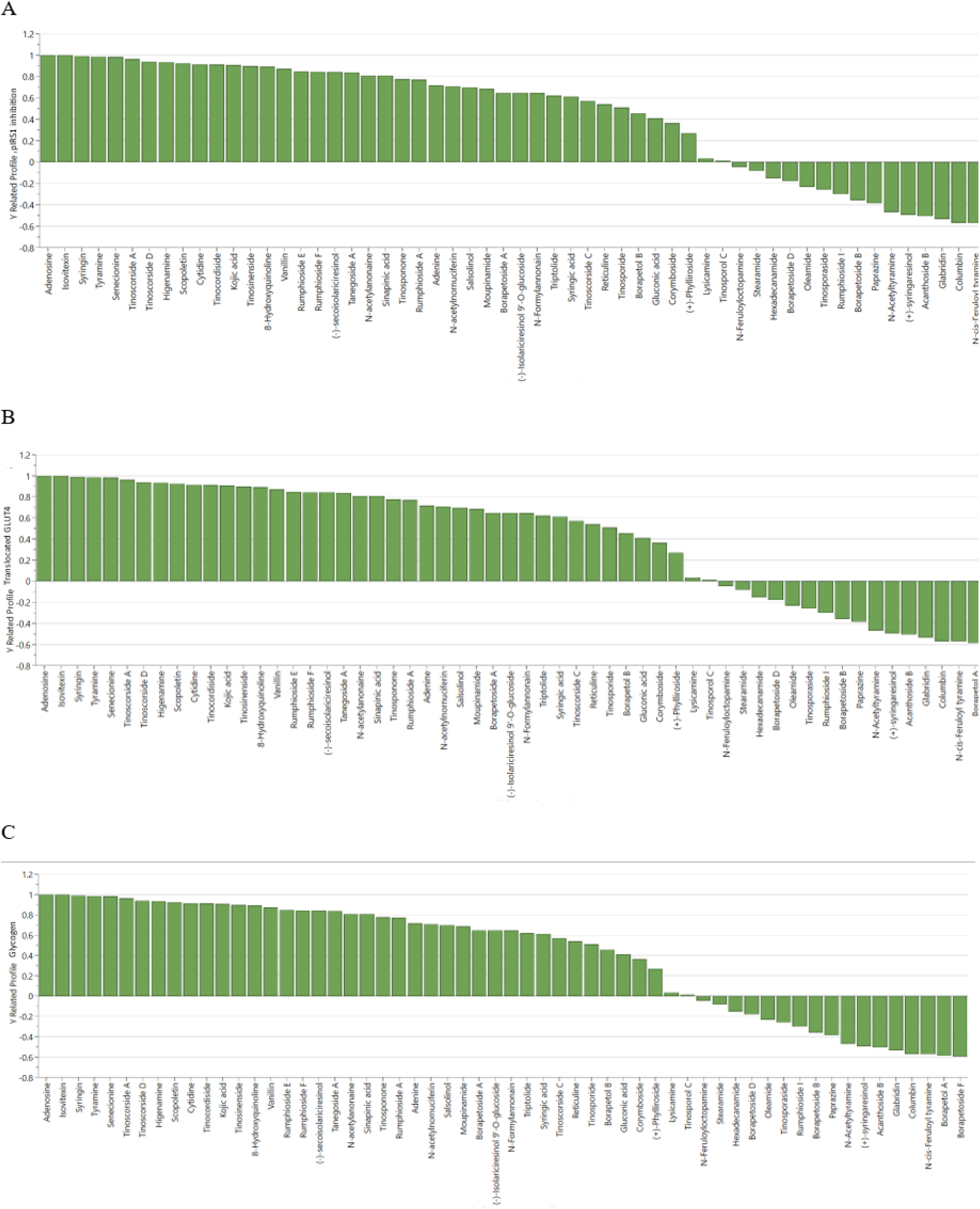
The OPLS Y-related profiles plots. (A) inhibition of pIRS1 ser312, (B) translocated GLUT4, and (C) Glycogen content. The X-axis represents the identified TCS metabolites, and the Y-axis represents the Y-related coefficients. The metabolites have been ranked based on the Y coefficient values, with the highest values on the left and the lowest values on the right.

### 3.4. Validation the TCS active metabolites using *in silico* molecular docking study

AutoDockTools was used to validate the accuracy of TCS active compound revealed by metabolomics study. This *in silico* study could determine the binding activity between primary active metabolites and GLUT4 as their key therapeutic target. The energy level denotes the binding strength, with lower energy levels indicating a stronger binding capacity. Typically, if the docking calculation score falls below −7 kcal/mol, the binding affinity is classified as potent (An et al., 2021). Based on the data presented in Table 3, the computed binding energies between ten primary active components to GLUT4 demonstrated a close or even lower value than −7. All the active metabolites have a higher binding affinity than metformin. The top five active metabolites consist of tinoscorside D, tinoscorside A, senecionin, higenamine, and tinocordiside suggesting a favorable binding interaction. The ligand binding site of the docking simulation result showed a remarkable active sites of GLUT4 as plotted in Figure 8 (Berman, 2000). The highest binding affinity towards the GLUT4 protein model was tinoscorside D, with a binding energy of −10.92 kcal/mol.

**Figure 8.**
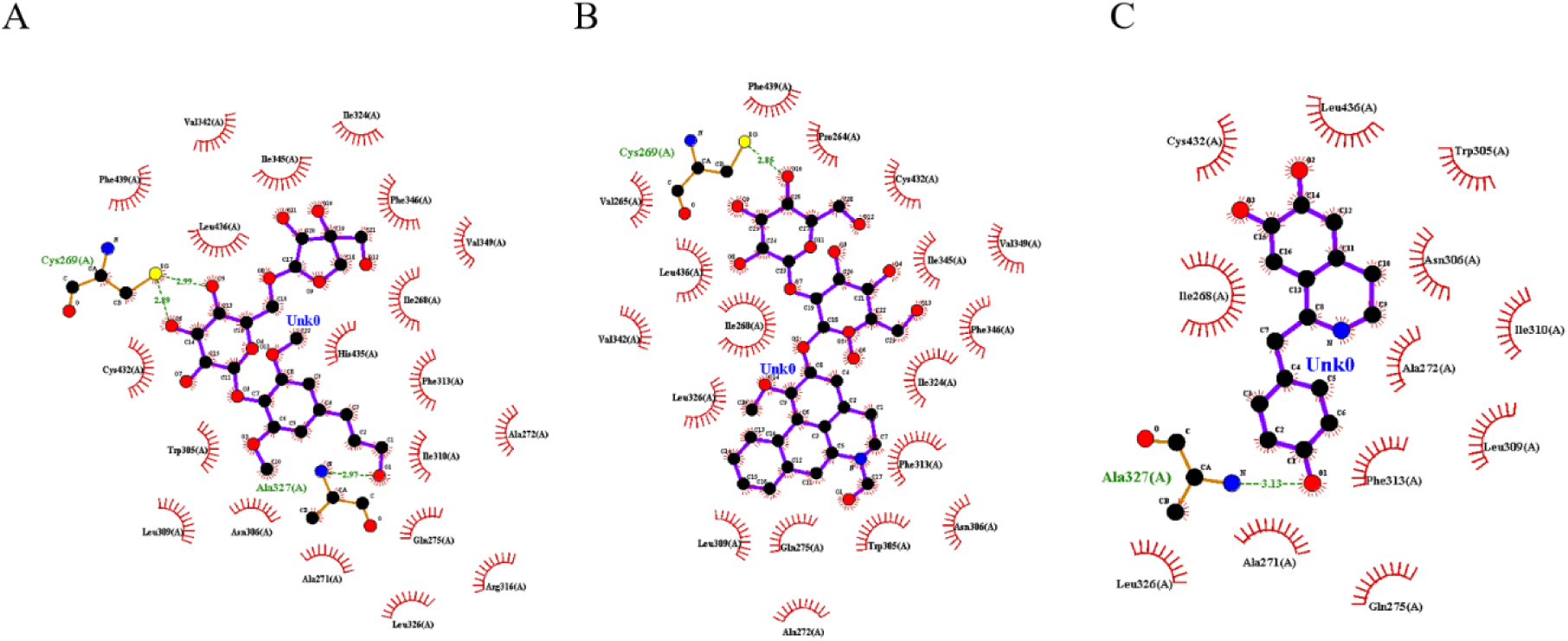
Ligand-protein docking plot. (A) tinoscorside D (B) tinoscorside A, and (C) higenamine to GLUT4. Black dots represent carbon atoms, red represents hydrophobic residues, blue represents nitrogen atoms, and yellow represents sulfur atoms. Dashed green lines represent hydrogen bonds, and red curved lines represent salt bridges, with the amino acids that form the bonds indicated.

**Table 3.** Virtual docking of ten active metabolites from TCS to GLUT4 protein model.

| No | Metabolites | $\Delta G$<br>(kcal/mol) | Inhibition<br>constant | Hydrogen bond |
| --- | --- | --- | --- | --- |
| 1 | Tinoscorside D | -10.92 | 9.81 nM | Cys269 |
| 2 | Native ligand LX81 | -10.64 | 15.85 nM | Ala327 |
| 8 | Tinoscorside A | -9.57 | 15.3 uM | Cys269 |
| 3 | Senesionin | -9.11 | 211.1 nM | Not exist |
| 4 | Higenamine | -9.11 | 208.36 nM | Ala327 |
| 5 | Tinocordiside | -8.53 | 555.13 nM | Val265, Cys269, Cys432 |
| 6 | Scopoletin | -6.7 | 12.29 uM | Arg316, Ala327 |
| 7 | Syringine | -6.61 | 14.22 uM | Arg316, Cys432 |
| 9 | Tyramine | -4.92 | 249.02 uM | Asn306, Ala327 |
| 10 | Cytidine | -4.48 | 524.44 uM | Asn306, |
| 11 | Isovitexin | -3.85 | 1.51 mM | Gln275, Cys432 |
| 12 | Metformin | -3.64 | 2.14 mM | Val265, Cys269, Cys432 |

The discovery of three active compounds in brotowali, namely tinoscorside D, higenamine, and tinoscorside A, is a novel finding that has important implications for the development of new treatments for insulin resistance. These findings are consistent with our previous study, which identified tinoscorside A as the active compound in brotowali (Zuhri et al., 2022). Tinoscorside A, despite being the least abundant compound in the active fraction, exhibits a high correlation with bioactivity, as evidenced by the area under the curve in the mass spectra. These results suggest that tinoscorside A may be a promising candidate for the development of new antidiabetic drugs (Tabel 1).

Tinoscorside D is a member of phenolic glycoside group that have previously been reported to be present in the plant *Tinospora cordifolia*. Our study identified the presence of this compound in *T. crispa*, a plant that belongs to the same genus and shares similarities in its chemical composition and bioactivity (Chi et al., 2016). Higenamine is an alkaloid compound that has also been reported to be present in *T. crispa* and is known for its active role as an antihypertensive agent (Praman et al., 2012).

These three compounds are minor compounds that are difficult to detect in natural product formulations. This is because natural medicinal sources are complex and require repeated fractionation and characterization, which can be time-consuming and expensive. Metabolomics methods are considered capable of expediting the identification process of both minor and major active compounds (Salem et al., 2020; Wishart, 2016; Yuliana et al., 2014).

The identification of active compounds in this study showed a different result from the previous research that reported borapetoside A and borapetoside C as the active compounds of *T. crispa* in insulin sensitization activity (Lam et al., 2012; Ruan et al., 2013). These previous studies used *in vitro* and *in vivo* methods to investigate the effects of these compounds, and they found that borapetoside A increased glycogen synthesis and decreased plasma glucose activity (Ruan et al., 2013). However, in this study, we used metabolomics to identify active compounds in TCS considering metabolomics as a powerful tool that can identify multiple compounds simultaneously. Our findings suggest that the three active compounds detected through metabolomics have great potential as candidate of new drugs for addressing insulin resistance.

## 4. Conclusion

The recent study deciphered, for the first time, tinoscorside D, tinoscorside A, and higenamine as the active constituents of TCS to insulin insulin resistance by combined *in vitro*, metabolomics, and *in silico* methods. Our data demonstrated that TCS could reduce IRS1 ser312 phosphorylation, promote GLUT4 translocation, and increase glycogen. TCS eventually ameliorated L6.C11 induced insulin resistance skeletal muscle cells through multi-component and multi-target effects. This valuable finding may provide theoretical support for further study as drug candidates of novel insulin sensitizing agents. However, our findings are preliminary and still need to be proven in *in vivo* studies and clinical trials as future research.

## List of abbreviations

T2DM: Type 2 Diabetes Mellitus
GLUT4: glucose transporter 4
*T. crispa*: *Tinospora crispa* (L.) Hook. f. & Thomson
pIRS: phosphorylated insulin receptor substrate
TCS: *Tinospora crispa* (L.) Hook. f. & Thomson stem;
MFI: Mean Fluorescence Intensity
DAPI 4’,6-diamidino-2-phenylindole: MS, mass spectrum
ppm: part per million
MVDA: multivariate data analysis
OPLS: Orthogonal Projection to Latent Square
PCA: Principal Component Analysis
RCSB: Research Collaboratory for Structural Bioinformatics
PDB: Protein Data Bank
SD: standard deviation
IR: insulin receptor
ser-312: serine 312
Met: metformin
VIP: Variable of Importance Person

## Author contributions

UMZ design and performed the study and drafted the manuscript; FF coordinated the technical support and funding, EHP designed the study and revised the manuscript; NDY designed and revised the manuscript; LE participated on the study and revised the manuscript; AC coordinated technical support and funding. All authors read and approved the manuscript.

