## Supplementary material 1, 2, 3, and 4 for "Exploration of the main active metabolites from *Tinospora crispa* (L.) Hook. f. & Thomson stem as insulin sensitizer in L6.C11 skeletal muscle cell by integrating *in vitro*, metabolomics, and molecular docking"

Supplementary material 1.

% Inhibition of pIRS1 ser312 treated with TCS fractions.

| Replication | % inhibition pIRS1 ser312 treated with TCS fraction | | | | | | | | | | | Metformin control |
|---|---|---|---|---|---|---|---|---|---|---|---|---|
| | F1 | F2 | F3 | F4 | F5 | F6 | F7 | F8 | F9 | F10 | F11 | |
| A | 38.73 | 39.36 | 42.34 | 42.99 | 34.95 | 25.97 | 44.30 | 27.51 | 30.94 | 31.92 | 5.79 | 74.60 |
| B | 38.15 | 36.02 | 49.99 | 27.65 | 34.27 | 34.84 | 46.66 | 23.69 | 39.72 | 25.93 | 7.76 | 70.69 |
| C | 45.64 | 21.56 | 44.71 | 33.37 | 28.71 | 34.47 | 41.95 | 19.31 | 38.77 | 15.00 | 11.23 | 85.27 |
| Mean | 40.84 | 32.31 | 45.68 | 34.67 | 32.64 | 31.76 | 44.30 | 23.50 | 36.48 | 24.28 | 8.26 | 76.85 |
| SD | 3.40 | 7.72 | 3.20 | 6.33 | 2.80 | 4.09 | 1.92 | 3.35 | 3.94 | 7.00 | 2.25 | 6.16 |

Supplementary material 2.

% Increasing of translocated GLUT4 treated with TCS fractions.

| Replication | % increasing of translocated GLUT4 treated with TCS fraction | | | | | | | | | | | Metformin control |
|---|---|---|---|---|---|---|---|---|---|---|---|---|
| | F1 | F2 | F3 | F4 | F5 | F6 | F7 | F8 | F9 | F10 | F11 | |
| A | 40.42% | 32.20% | 75.80% | 37.09% | 30.58% | 21.30% | 44.61% | 19.94% | 46.55% | 25.20% | 24.10% | 34.31% |
| B | 40.13% | 29.14% | 70.25% | 39.69% | 32.64% | 40.13% | 45.95% | 23.36% | 50.59% | 35.60% | 20.55% | 40.02% |
| C | 45.15% | 34.36% | 81.08% | 38.44% | 32.07% | 21.85% | 51.53% | 20.45% | 46.87% | 31.21% | 27.29% | 40.36% |
| Mean | 41.90% | 31.90% | 75.71% | 38.41% | 31.76% | 22.85% | 47.36% | 21.25% | 48.00% | 30.67% | 23.98% | 38.23% |
| SD | 2.30% | 2.14% | 4.42% | 1.06% | 0.87% | 2.30% | 3.00% | 1.50% | 1.83% | 4.27% | 2.75% | 2.77% |

Supplementary material 3.

Glycogen content in L6.C11 skeletal muscle cells treated with TCS fractions.

| Replication | % glycogen content increasing treated with TCS fraction | | | | | | | | | | |
|---|---|---|---|---|---|---|---|---|---|---|---|
| | F1 | F2 | F3 | F4 | F5 | F6 | F7 | F8 | F9 | F10 | F11 |
| A | 32.88% | 41.49% | 59.56% | 62.88% | 64.24% | 72.31% | 45.70% | 66.99% | 36.11% | 32.54% | 52.76% |
| B | 36.34% | 40.75% | 69.84% | 48.80% | 56.91% | 56.73% | 52.88% | 47.35% | 66.83% | 46.55% | 32.46% |
| C | 41.55% | 43.60% | 61.26% | 60.16% | 50.54% | 54.86% | 50.29% | 64.49% | 46.99% | 55.31% | 47.71% |
| Mean | 36.92% | 41.95% | 63.55% | 57.28% | 57.23% | 61.30% | 49.63% | 59.61% | 49.98% | 44.80% | 44.31% |
| SD | 4% | 1% | 4% | 6% | 6% | 8% | 3% | 9% | 13% | 9% | 9% |

Supplementary material 4.

Representative confocal images of L6.C11 treated TCS fractions and control groups

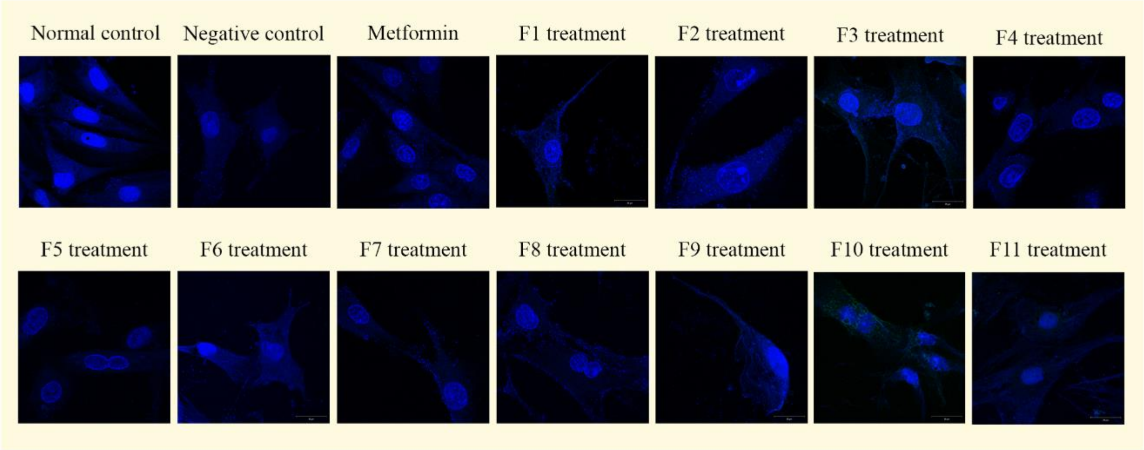


Representative images of laser confocal imaging for GLUT4 quantification at 63x magnification from all experimental groups. The F1-F11 group represents insulin-resistant cells treated with F3 at 400 ppm, the metformin group was treated with metformin, while the negative group remains untreated. The normal group consists of uninduced cells treated only with the solvent used in F3 (0.5% (v/v) DMSO in DMEM). Images were captured at 490 nm for GLUT4 imaging and 510 nm for DAPI imaging.
